# Proteomics- and BRET-screens identify SPRY2 as Ras effector that impacts its membrane organisation

**DOI:** 10.1101/2025.06.13.659437

**Authors:** Karolina Pavic, Fiona Elizabeth Hood, Carla Jane Duval, Ganesh babu Manoharan, Christina Laurini, Farid Ahmad Siddiqui, Stephanie P Mo, Ian Andrew Prior, Daniel Kwaku Abankwa

## Abstract

K-Ras functions within nanoscale proteo-lipid domains of the plasma membrane, but few regulators of its membrane organisation are known. We combined TurboID-based proximity proteomics with a secondary BRET screen to identify eight novel K-Ras G-domain interactors. We focused on APLP2 and SPRY2 for further characterisation. APLP2 binds K-Ras indirectly via C-Raf, while SPRY2 exhibits properties of a novel effector. Co-immunoprecipitation and BRET assays revealed that the SPRY2 C-terminal fragment (residues 161–315) binds oncogenic RasG12V more strongly than the full-length protein. Both forms localise to the plasma membrane, but this localisation and binding to K-Ras is disrupted by inhibitors of K-Ras membrane anchorage or activity. Mutations at the predicted interface of K-Ras and SPRY2’s C-terminal region affect the interaction. Both full-length SPRY2 and its C-terminal fragment promote differentiation of C2C12 muscle cells, a process requiring MAPK pathway inhibition. Finally, SPRY2 also forms homo- or hetero-oligomers with SPRY4. We propose that active K-Ras recruits SPRY2 dimers to the membrane, where they bind Ras and block effector access.

## Introduction

Ras signalling plays a critical role in cell proliferation and differentiation and contributes to a wide range of cellular functions and phenotypes important for health and disease (Simanshu *et al*, 2017). The Ras protein operates like a molecular switch at the inside of the plasma membrane where it links growth factor receptor inputs to effector outputs. Ras effectors bind to Ras in a nucleotide-dependent manner, and their concentration at the plasma membrane leads to functional interactions that influence cell signalling networks (Smith, 2023). For instance, Raf family members are canonical Ras effectors that initiate the MAP kinase cascade, which is classically associated with cell proliferation.

The Ras family consists of four isoforms, H-Ras, N-Ras, K-Ras4A and K-Ras4B (hereafter referred to as K-Ras), that are structurally highly related and share a common set of regulators and effectors (Hobbs *et al*, 2016). While their G-domain is 80-90 % identical, the C-terminal hypervariable region (HVR) distinguishes them. The HVR of all Ras proteins becomes farnesylated thus anchoring Ras to cellular membranes. However, isoform-specific functional differences have been observed and in recent years this is thought to at least be partly due to isoform-specific differences in subcellular localisation and occupation of a mosaic of different cell surface signalling nanoclusters (Abankwa & Gorfe, 2020; Pavic *et al*, 2022).

The mechanistic links between nanoscale spatial organisation and Ras isoform functional specificity remains poorly understood because of the challenges in characterising the protein and lipid compositions of ephemeral signalling domains. One strategy for understanding the cellular context that Ras proteins operate in has been to employ proteomic analysis of the Ras interactome. Up to 210 proteins were detected as potential direct or indirect Ras interactors capable of being co-immunoprecipitated (Goldfinger *et al*, 2007). Several groups have since utilised BirA proximity-dependent biotin identification (BioID), able to detect not only direct interactors but also proteins in proximity to Ras (Roux *et al*, 2012). BirA fused to the Ras N-terminus catalyses the conjugation of biotin to proteins in ∼ 10 nm vicinity that can then be identified via streptavidin enrichment and proteomic analysis. Ras proximal proteome studies to date have typically relied on full length wild type and constitutively active Ras mutants (Adhikari & Counter, 2018; Beganton *et al*, 2020; Cheng *et al*, 2021; Kovalski *et al*, 2019). One of the challenges with this approach is the promiscuity of biotinylation since the proximal proteome of the full cellular itinerary of Ras from initial expression to all its destinations is labelled. Combination with a secondary orthogonal screen has been an effective way for prioritising hits (Adhikari & Counter, 2018). Another refinement has been TurboID that employs a mutated BirA optimised to significantly reduce the biotinylation timeline from 18-24 hours to 10 minutes (Branon *et al*, 2018).

In this study we aim to identify novel proteins that regulate K-Ras membrane organisation and function in dependence of its activation state.

## Results

### TurboID identifies the proximal proteome associated with the G-domain and mutationally activated K-Ras

We employed TurboID to identify proteins that operate in close proximity to K-Ras. Experiments were conducted to first identify the K-Ras proximal proteome and then to identify the dependence of this proximity on the K-Ras activation state (**Fig. 1A**). Stable isotope labelling by amino acids in cell culture (SILAC) was used to allow for ratiometric proteomic comparison of streptavidin enrichment between each set of comparators (Ong *et al*, 2002). SILAC uses heavy isotope variants of arginine (R) or lysine (K) that allows peptides isolated from samples grown with these isotopes to be distinguished in the mass spectrometer. The available combinations of arginine and lysine isotopes mean that three experimental conditions (triplexes) can be compared in each mass spectrometry run. In all comparator triplexes, a TurboID + biotin control was included to allow pan-experiment comparison of all conditions. For proximity analysis, the other two comparators in a triplex were each of the K-Ras variants +/-biotin, whereas for activation dependence the comparators consisted of treatment with or without K-RasG12C-inhibitor (**G12Ci**), + biotin (**Figs. S1 and S2**).

**Figure 1.**
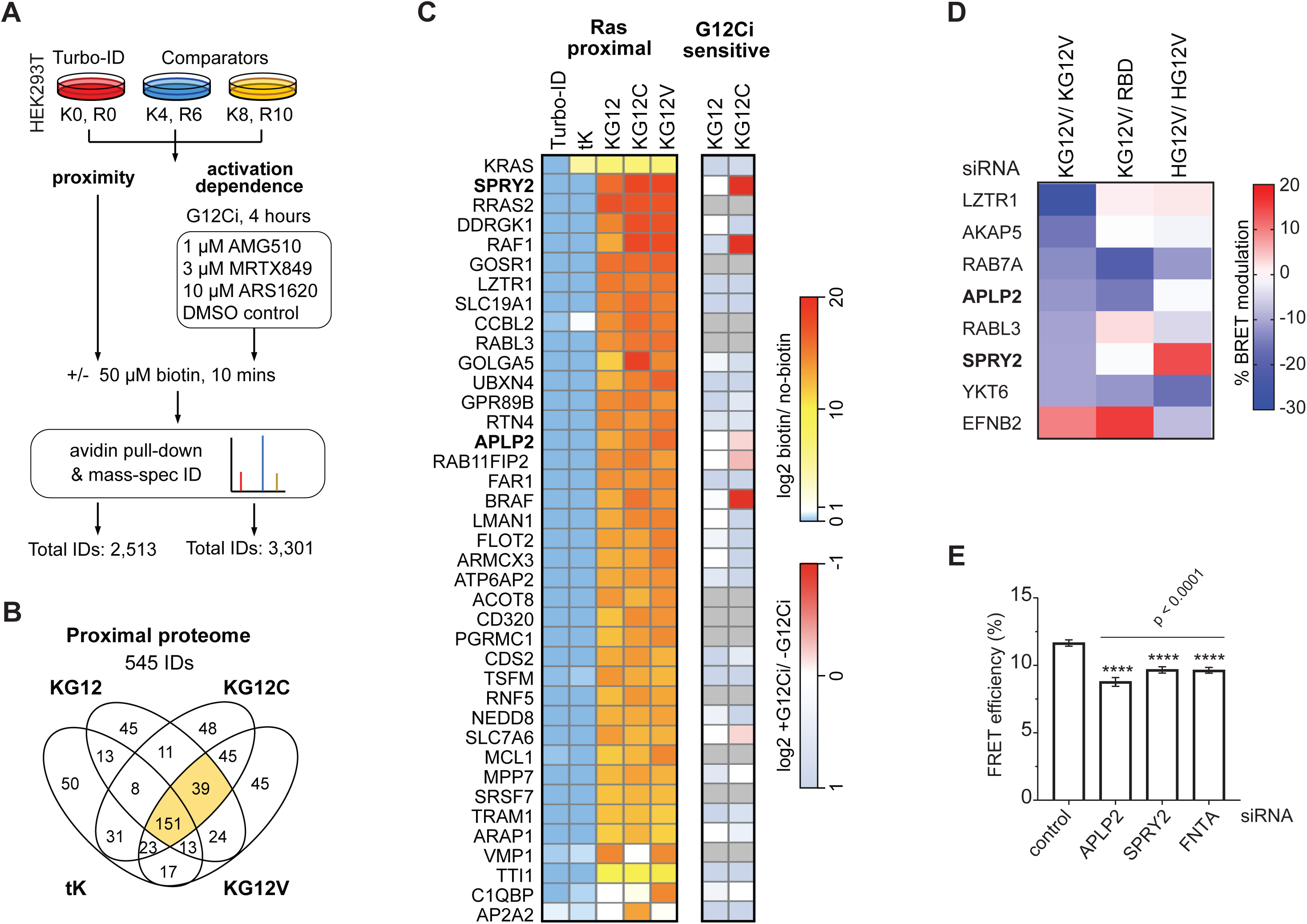
TurboID- and BRET-screening identifies bone fide K-Ras G-domain interactors that modulate its membrane organisation. (**A**) Schematic of the two TurboID experiments that detected proximity and activation dependent proximity to K-Ras variants. SILAC labelling employs isotopes of arginine (R0, 6, 10) and lysine (K0, 4, 8) to allow mass-spectrometry identification and ratiometric comparison of relative protein biotinylation in each condition following streptavidin enrichment. (**B**) Venn Diagram by TurboID-probes, wt K-Ras (KG12), K-RasG12D (KG12C), K-RasG12V (KG12V) and tK. Total IDs were all proteins detected in at least one condition across N = 3 experiments. The proximal proteome shortlist represents all hits that were ≥ 2-fold enriched versus the equivalent non-biotinylated control condition and also not proximal to the TurboID control. Data are rank order from top to bottom of most to least enriched across all three variants. (**C**) Relative enrichment of the 39 proximal hits observed in all three of the full-length K-Ras variant conditions together with the activation dependence of their proximity. (**D**) Modulation of the BRET signal after siRNA-mediated knockdown of indicated TurboID hits in BRET-assays to measure K-RasG12V membrane organisation (KG12V/ KG12V), K-RasG12V/ C-Raf-RBD interaction (KG12V/ RBD) and H-RasG12V membrane organisation (HG12V/ HG12V) in HEK cells from N = 4 biological repeats. (**E**) Quantification of FLIM-FRET imaging data of HEK293T cells transfected with pmEGFP-K-RasG12V/ pmCherry-K-RasG12V after siRNA-mediated knockdown of indicated genes of interest. Means ± SEM of N = 3 biological repeats are plotted. For APLP2 siRNA, N = 2 biological repeats. Per condition, 50-100 cells were analysed. Statistical comparisons were done using one-way ANOVA and differences to the scrambled siRNA control are indicated.

For K-Ras proximity, we compared three full-length K-Ras variants comprising wild type and the G12C and G12V activated mutants of K-Ras that are commonly found in cancer patients (Prior *et al*, 2020). We also included the K-Ras fragment tK, which comprises residues 175-188 of the HVR (Prior *et al*, 2001). While it is targeted to the plasma membrane, it lacks the G-domain (residues 1-165) that is necessary for most, if not all, currently known Ras interactions (Prior *et al*., 2001). Three biological replicates of each experiment were conducted, and the total number of proteins seen in each condition for the proximal proteome experiment was 545 (**Fig. 1B, Data S1)**. Shortlisting was based on observing an average of ≥ 2-fold enrichment over the control without biotin across the replicates together with also not being proximal to the cytoplasmic TurboID control (≥ 2-fold + biotin vs – biotin TurboID). Thus, 190 proteins were identified across all K-Ras variants, with 39 proteins observed just with the full-length variants but not tK suggesting their proximity depends on the G-domain (**Fig. 1C**).

For activation-dependence of K-Ras proximity, we tested three different selective G12Ci with K-RasG12C and K-Ras wild type conditions. Direct covalent inhibition of mutant K-Ras is an improvement over previous strategies of comparing wild type with constitutively active K-Ras mutants. The proximity of 132 proteins to K-RasG12C was sensitive to G12Ci (**Data S1**) and comprised canonical Ras effectors such as Raf and PI3-kinase family members. Raf1 (C-Raf), B-Raf and SPRY2 showed a prominent G12Ci-sensitivity of their K-RasG12C proximity labelling (**Fig. 1C, Figs. S1 and S2**). This might be expected for the former two, given that Raf proteins are well-established Ras effectors. Yet for SPRY2 the G12Ci-sensitive proximity suggests a potentially novel, activation state-dependent interaction.

### Profiling for Ras-membrane organisation modulators by BRET

To functionally profile proteomic hit proteins as impacting on K-Ras membrane organisation, we performed bioluminescence resonance energy transfer (BRET) experiments following knock-down of the hit genes using siRNA.

Our primary BRET-screen employed our well-established K-RasG12V-membrane organisation BRET-biosensor, consisting of the BRET-pair RLuc8-K-RasG12V and GFP2-K-RasG12V expressed in HEK293-EBNA cells (**hereafter HEK**). An analogous FRET-biosensor reported on the loss of K-Ras-nanoclustering, -membrane anchorage and -lipidation with a drop in the FRET-signal (Parkkola *et al*, 2021). In agreement with this, we previously demonstrated that the K-RasG12V BRET-biosensor responds to an inhibition of Ras trafficking chaperones and Ras lipidation (Duval *et al*, 2024; Kaya *et al*, 2024; Manoharan *et al*, 2023; Okutachi *et al*, 2021).

Validation of this assay in the screening setting revealed that ablation of FNTA, the gene encoding for the common ⍺-subunit of farnesyl-and geranylgeranyl-transferases led to the strongest drop of the BRET-signal (**Fig. S3A**). A clear effect was also observed after knockdown of trafficking chaperones (PDE6D, CALM1), while nanocluster modulators (LGALS3, TP53BP2) had a less pronounced effect (**Fig. S3A**). This latter observation contrasts to our previous FRET-measurements, which displayed a much larger dynamic range (Manoharan *et al*, 2022; Posada *et al*, 2016; Siddiqui *et al*, 2020; Siddiqui *et al*, 2021).

Based on these validation data, we set a 10 %-change in the BRET-signal as a threshold for high confidence BRET-hits of this primary BRET-screen. A selection of 29 hits from the putative G-domain binder hit list and a meta-analysis of four other K-Ras-proximal proteome studies were then screened (**Fig. S3B**). The meta-analysis identified 1,366 proteins proximal to at least one isoform of Ras (Adhikari & Counter, 2018; Beganton *et al*., 2020; Cheng *et al*., 2021; Kovalski *et al*., 2019). However, only 12 K-Ras proximal proteins were common to all four other studies and included Raf isoforms and the Ras inactivator NF1 (**Data S1**). From those proteins that were common to three or four of these studies, we selected a subset of 21 for further investigations.

While 4 of 8 (50 %) of the G-domain binder list were confirmed as BRET hits, only 4 of 21 (∼ 20 %) from the meta-analysis list qualified as such, suggesting that our differential TurboID-screening approach is advantageous to identify putative K-RasG12V membrane organisation modulators (**Fig. 1D, Fig. S3B**). All of the BRET-screen hits reduced K-RasG12V membrane organisation BRET after their siRNA-mediated depletion, except for EFNB2, which increased it (**Fig. 1D**).

To functionally profile hits further in secondary BRET-screens, we employed two additional biosensors. Using the RLuc8-K-RasG12V and C-Raf-RBD-GFP2 biosensor, we could detect perturbations of Ras effector binding. This included also loss of Ras nanoclustering, which we previously showed reduces the recruitment efficiency of effectors from the cytosol (Guzman *et al*, 2014; Steffen *et al*, 2024) (**Fig. 1D, Fig. S3C**). The third BRET-assay served to assess relative K-Ras selectivity by detecting H-Ras membrane organisation using RLuc8-H-RasG12V and GFP2-H-RasG12V as biosensor (**Fig. 1D, Fig. S3C**).

Based on this characterisation, we decided to focus on two genes from the BRET-screen hits, APLP2 and SPRY2. APLP2 is not known for any function in concert with K-Ras. SPRY2 belongs to the family of four SPRY-or sprouty-proteins, which are frequently observed hits in Ras proximal proteome screens (Adhikari & Counter, 2018; Beganton *et al*., 2020; Cheng *et al*., 2021; Kovalski *et al*., 2019). To confirm our BRET-screen results for these proteins, we used the more sensitive fluorescence lifetime imaging microscopy (FLIM)-Förster resonance energy transfer (FRET) approach. FLIM-FRET data supported a highly significant reduction of K-RasG12V membrane-FRET upon knockdown of either APLP2 or SPRY2 (**Fig. 1E**). Given that it is unknown how these proteins bind to Ras and impact on its membrane organisation, we continued their investigation.

### APLP2 interacts with C-Raf and oncogenic K-Ras

APLP2 (Amyloid precursor-like protein 2) is a member of the evolutionary conserved amyloid-precursor-like (APP) protein family. APLP2 is a ubiquitously expressed trans-membrane glycoprotein that becomes proteolytically activated by secretases (Pandey *et al*, 2016). APLP2 is aberrantly expressed in different types of cancers, with over-expression as well as proteolytic cleavage positively correlating with cell growth and migration (Pandey *et al*., 2016). In the genetically engineered KRAS-G12D-driven KPC-mouse model for spontaneous pancreatic cancer, deletion of APLP2 significantly prolonged survival and reduced metastasis (Poelaert *et al*, 2021). Given this genetic interaction of K-Ras and APLP2, we wanted to validate the biochemical interaction of these two proteins.

Using co-immunoprecipitation experiments, we characterised the engagement of APLP2 with Ras and Raf proteins. Endogenous APLP2 was ∼15-fold more co-immunoprecipitated with GFP2-tagged C-Raf than with GFP2-tagged K-RasG12V (**Fig. 2A,B**). This was in line with the BRET-profiling data, showing that depletion of APLP2 affects both the K-RasG12V/ K-RasG12V-and K-RasG12V/ C-Raf-RBD-BRET signal (**Fig. 1D**).

**Figure 2.**
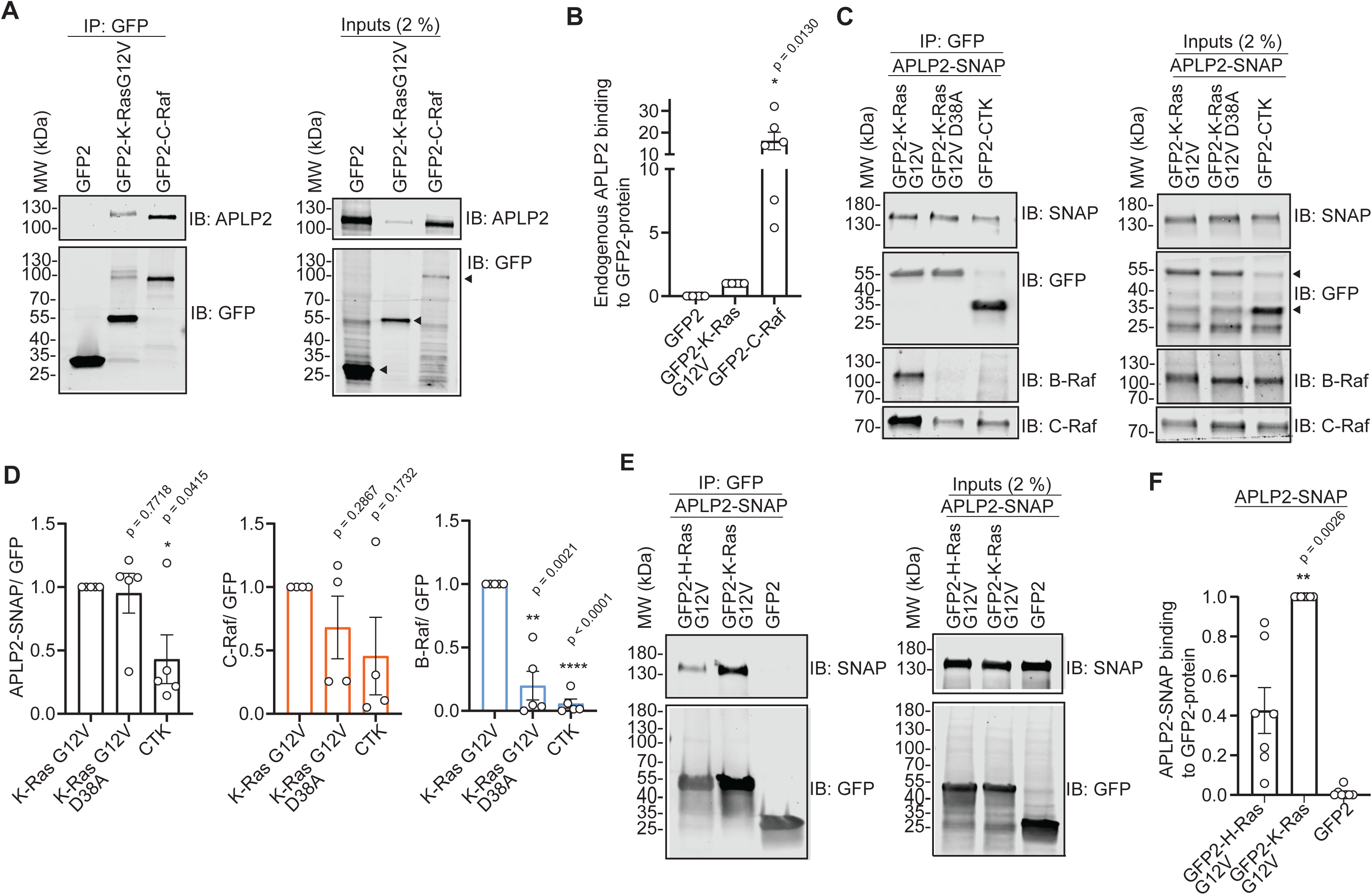
APLP2 preferentially interacts with C-Raf and less with K-RasG12V. (**A,B**) Representative blots of GFP-Trap pull-downs (left) of endogenous APLP2 from HEK lysates (right) using GFP2-K-RasG12V or GFP2-C-Raf (A) with quantified means ± SEM of N = 6 or 7 biological repeats of pull-down data (B). Arrowheads mark specific bands of expressed constructs. (**C,D**) Representative blots of GFP-Trap pull-downs (left) of APLP2-SNAP and endogenous C-Raf and B-Raf in the presence of the former from HEK lysates (right) using GFP2-K-Ras derived constructs as indicated with GFP2-CTK serving as control (C) with quantified means ± SEM of N ≥ 4 biological repeats of pull-down data (D). Arrowheads mark specific bands of expressed constructs. (**E,F**) Representative blots of GFP-Trap pull-downs (left) of APLP2-SNAP from HEK lysates (right) using GFP2-K-RasG12V, GFP2-H-RasG12V or GFP2 (control) as indicated (E) with quantified means ± SEM of N = 7 biological repeats of pull-down data (F).

To understand if the K-Ras interaction was sensitive to the D38A-mutation, which abrogates interactions with all major effectors (Akasaka *et al*, 1996), we performed GFP-Trap pull-down experiments of C-terminally SNAP-tagged full length APLP2 with GFP2-K-Ras constructs from HEK cell lysates. All signal obtained with GFP2-CTK, which encodes only the membrane anchoring HVR sequence (residues 166-188) of K-Ras, was considered background.

While both endogenous C-Raf and B-Raf co-immunoprecipitated more with K-RasG12V than K-RasG12V-D38A, APLP2-SNAP co-immunoprecipitated to about the same extent with both K-Ras mutants (**Fig. 2C,D**). This is in line with APLP2 primarily engaging with C-Raf (**Fig. 2A,B**). BRET-screening data suggested a specific effect of APLP2 on K-RasG12V-but not on H-RasG12V-membrane organisation (**Fig. 1D**). In line with this GFP2-K-RasG12V co-immunoprecipitated >2-fold more APLP2-SNAP than GFP2-H-RasG12V (**Fig. 2E,F**).

In conclusion, both BRET-validation and co-immunoprecipitation data suggest that APLP2 may only indirectly engage with active K-RasG12V via Raf-proteins, and less with H-RasG12V. APLP2 may thus stabilise Ras/ Raf-complexation at the plasma membrane and (Raf-dependent) K-Ras nanoclustering, consistent with our previous model for active nanocluster (Blazevits *et al*, 2016; Siddiqui *et al*., 2021; Steffen *et al*., 2024).

### SPRY2 interacts with oncogenic K-Ras predominantly through its C-terminal region

SPRY2 is ubiquitously expressed and the evolutionary most conserved member of the four human or mouse SPRY proteins (Guy *et al*, 2009; Puranik *et al*, 2024). SPRY proteins are characterised by their C-terminal cysteine-rich domain (CRD), which contains several palmitoylation sites that are important for their localisation to the plasma membrane (Lim *et al*, 2002; Locatelli *et al*, 2020). The N-terminus of SPRYs is more diverse and features a c-Cbl-tyrosine kinase-binding (Cbl-TKB) motif followed by a serine-rich motif (SRM). At the plasma membrane SPRY2 is phosphorylated on a conserved Tyr-55 in the Cbl-TKB motif, which is associated with ubiquitin mediated regulation of SPRY stability (Guy *et al*., 2009).

Drosophila SPRY was identified as an antagonist of the Ras-MAPK pathway during fly trachea development (Hacohen *et al*, 1998). This ability to down-regulate Ras-signalling in response to a variety of growth factor stimuli is preserved in vertebrates (Hanafusa *et al*, 2002). It was previously proposed that phosphorylated SPRY negatively regulates Ras-MAPK-signalling by sequestering the adaptor protein Grb2, thus preventing activation of SOS (Hanafusa *et al*., 2002). Others suggested SPRY proteins to inhibit the MAPK-but not PI3K-pathway downstream of Grb2-SOS (Gross *et al*, 2001). However, several biochemical and proteomics studies identified SPRY2 as a potential K-Ras-(Adhikari & Counter, 2018; Go *et al*, 2021; Kovalski *et al*., 2019) or H-Ras-interactor (Lito *et al*, 2008).

We therefore wanted to identify the structural determinants for a potentially direct interaction between K-Ras and SPRY2. Using co-immunoprecipitation and cellular BRET-interaction studies, we verified our BRET-screening data, which suggested an interaction preference for K-Ras but no impact of SPRY2-depletion on effector engagement (**Fig. 1D**). Indeed, GFP-Trap pull-down confirmed that endogenous SPRY2 from HEK cell lysates binds ∼ 2.5-fold more to GFP2-K-RasG12V than GFP2-C-Raf (**Fig. 3A,B**).

**Figure 3.**
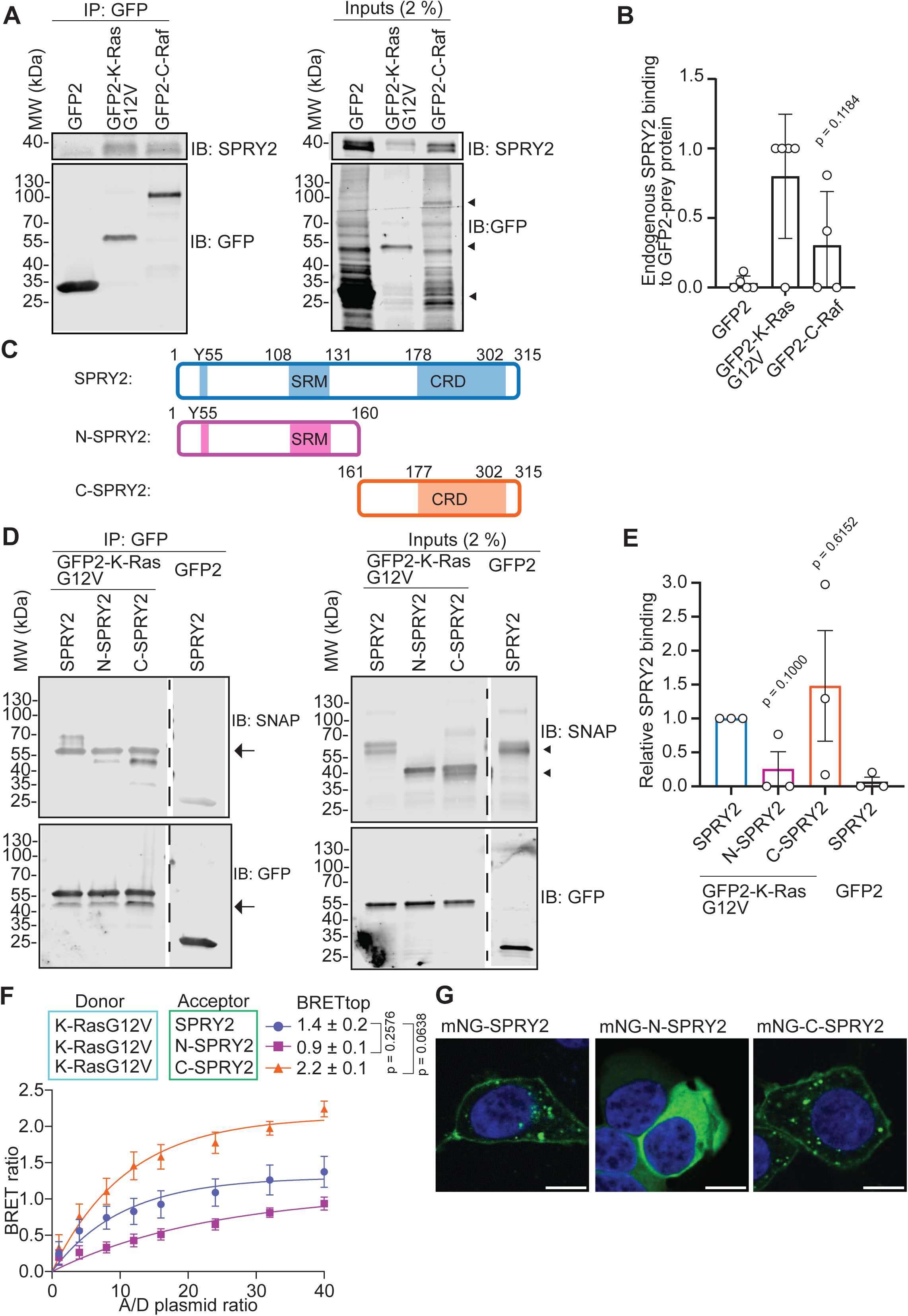
SPRY2 interacts with K-Ras predominantly via its C-terminal half. (**A,B**) Representative blots of GFP-Trap pull-downs (left) of endogenous SPRY2 from HEK lysates (right) using GFP2-K-RasG12V or GFP2-C-Raf (A) with quantified means ± SEM of N = 5 biological repeats of pull-down data (B). Arrowheads mark specific bands of expressed constructs. (**C**) Schematic of the SPRY2-fragment constructs used in pull-down and BRET experiments. SRM indicates serine rich motif and CRD Cysteine rich domain. The Y55 marks the tyrosine of the Cbl-TKB binding motif. (**D,E**) Representative blots of GFP-Trap pull-downs (left) using indicated SNAP-tagged SPRY2-constructs from lysates of HEK (right) expressing GFP2-K-RasG12V or GFP2 (control) (D) with quantified means ± SEM of N = 3 biological repeats of pull-down data (E). Arrowheads mark specific bands of expressed constructs. (**F**) BRET-titration curves of the nanoLuc-K-RasG12V interaction with mNG-tagged SPRY2 fragments as introduced in (C) acquired in HEK cells. N=3 biological repeats. (**G**) Confocal imaging showing subcellular localization of mNG-tagged SPRY2-constructs in HEK cells. Scale bar = 10 µm.

To identify which part of SPRY2 interacts with K-RasG12V, we studied the interactions of SNAP-tagged N-terminal SRM-comprising (N-SPRY, residues 1-160) and C-terminal CRD-comprising fragments (C-SPRY2, residues 161-315) (**Fig. 3C**). As compared to the full-length protein, N-SPRY2 showed a ∼ 5-fold reduced interaction with K-RasG12V, while C-SPRY2 interaction was ∼ 1.5-fold increased (**Fig. 3D,E**). This was confirmed by BRET-interaction data between mNeonGreen (mNG)-tagged SPRY2-fragments and nanoLuc (nL)-tagged K-RasG12V. The BRET-signal was 1.6-fold reduced with N-SPRY2, while that of C-SPRY2 was 1.6-fold increased (**Fig. 3F**). The same order of BRET-interactions was seen with H-RasG12V (**Fig. S4A**).

High BRET-levels can emerge from co-localisation at the plasma membrane. We therefore analysed the distribution of the mNG-tagged SPRY2-constructs in HEK cells by confocal imaging. When observed under normal serum conditions, full length SPRY2 was found at the plasma membrane both in the presence (**Fig. S4B**) and absence of co-expressed mCherry-K-RasG12V (**Fig. 3G**). Also, C-SPRY2 localised in this way, while N-SPRY2 was predominantly in the cytoplasm (**Fig. 3G**). This distribution pattern of the constructs can be explained by the fact that the C-terminal CRD of SPRY2 contains 26 cysteine residues, some of which are known to be palmitoylated. Palmitoylation is important for targeting SPRY2 to the plasma membrane, and mutation of two cysteine residues, C265 and C268, re-distributes SPRY2 to the cytoplasm (Locatelli *et al*., 2020).

Thus, the C-terminal half of SPRY2 is sufficient to translocate it to the plasma membrane, where it can engage with RasG12V.

### SPRY2 recruitment to the plasma membrane is sensitive to lipidation and K-RasG12C inhibitors

We next wanted to understand if SPRY2 plasma membrane localisation depends on prenylated and active K-Ras. We therefore first treated cells with mevastatin, which blocks the synthesis of prenyl-pyrophosphate in the mevalonate pathway and thus Ras prenylation and membrane organisation associated BRET (Okutachi *et al*., 2021). Given that C-terminal palmitoylation of SPRY2 is required for its membrane anchorage, we reasoned that 2-bromopalmitate (2-BP) would directly block SPRY2 plasma membrane binding (Hedberg *et al*, 2011; Locatelli *et al*., 2020).

We first confirmed that nanoclustering-associated BRET of farnesylated K-RasG12V was significantly abrogated by mevastatin and that of palmitoylated and farnesylated H-RasG12V was sensitive to both mevastatin and/ or 2-BP (**Fig. S5A,B**).

Confocal imaging data suggested that mevastatin-treatment led to the emergence of a cytoplasmic pool of SPRY2, similar to that observed after the treatment with 2-BP. Combined treatment with mevastatin and 2-BP most efficiently redistributed SPRY2 to the cytoplasm (**Fig. 4A**). We then used BRET between K-RasG12V and SPRY2-fragments to quantify the impact of the lipidation inhibitors on their co-localisation-dependent interaction. If both proteins interact at the membrane, high BRET is expected, also due to the up-concentration at the membrane (Parkkola *et al*., 2021). However, if one or both cannot localise to the membrane due to blocked lipidation the BRET-signal would be lowered. BRET of full length mNG-SPRY2 with nL-K-RasG12V was significantly reduced after mevastatin treatment, but even more so after 2-BP incubation or treatment with both lipidation inhibitors (**Fig. 4B**). While the N-SPRY2 fragment localised predominantly to the cytoplasm (**Fig. 3G**), its BRET with K-RasG12V was still sensitive to mevastatin, suggesting that membrane localised K-Ras can still engage with N-SPRY2. Consistent with N-SPRY2 being devoid of the C-terminal palmitoylation sites, 2-BP did not affect the BRET, while the combination of mevastatin and 2-BP reduced the BRET to the same extent as mevastatin alone (**Fig. 4C**). C-SPRY2 showed the highest BRET with K-RasG12V, which was significantly reduced after mevastatin treatment and even more after 2-BP treatment, while the combination of both inhibitors could not further reduce the BRET (**Fig. 4D**).

**Figure 4.**
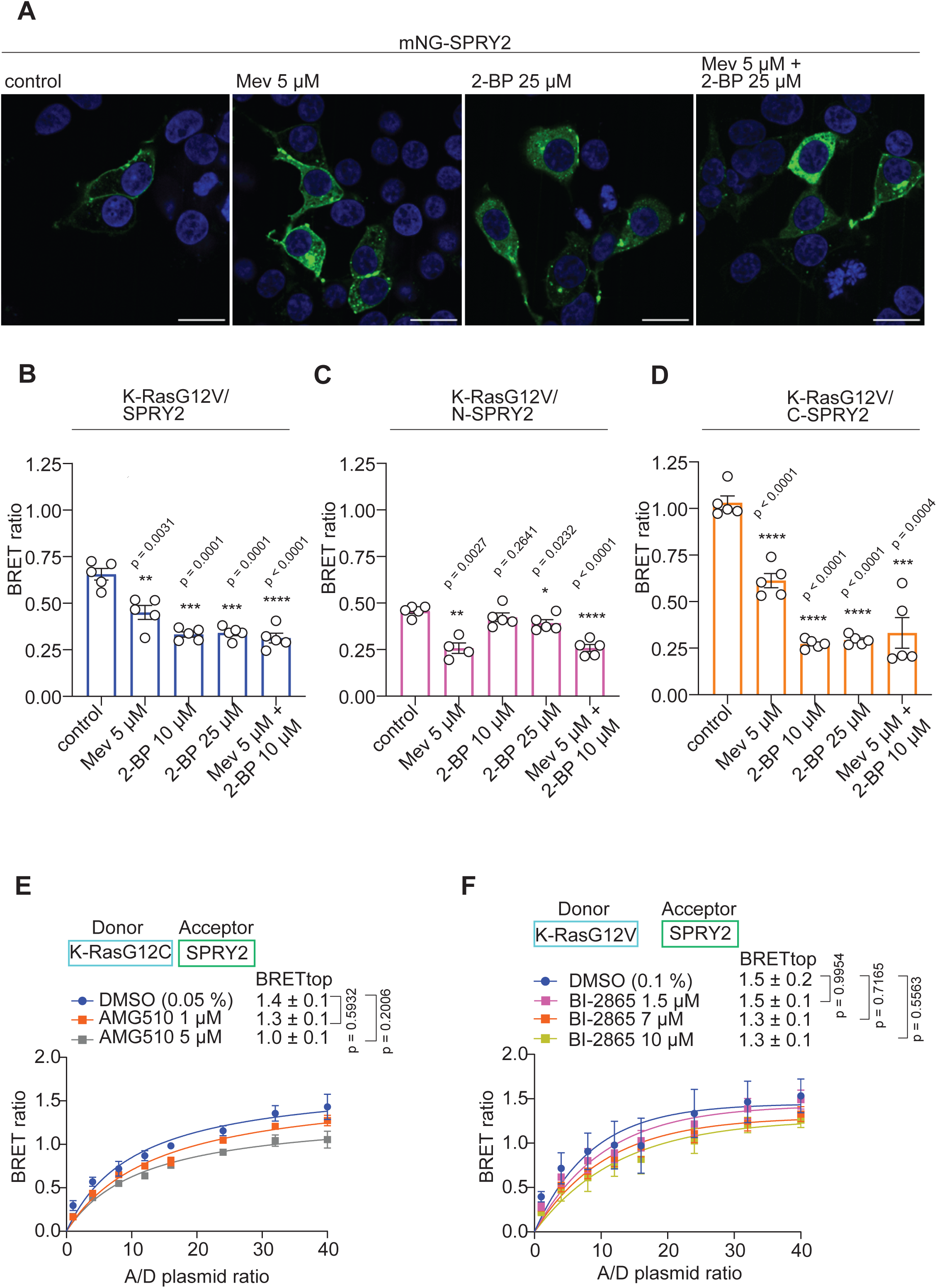
K-RasG12V engagement with SPRY2 is sensitive to lipidation inhibitors. (**A**) Confocal imaging showing that plasma membrane localisation of mNG-tagged SPRY2 in HEK cells is sensitive to lipidation inhibitors mevastatin (Mev) and/ or 2-bromopalmitate (2-BP). Scale bar = 20 µm. (**B-D**) Effect of mevastatin (Mev) and/ or 2-bromopalmitate (2-BP) on interaction-BRET of nanoLuc-K-RasG12V and mNG-SPRY2 (B), mNG-N-SPRY2 (C) and mNG-C-SPRY2 (D) in HEK cells (donor:acceptor plasmid ratio = 1:8). N = 5 independent biological repeats. (**E,F**) BRET-titration curves of the nanoLuc-K-RasG12V/ mNG-SPRY2 interaction after treatment with G12Ci AMG510 (E) or pan-K-Ras inhibitor BI-2865 (F) in HEK cells from N = 3 to 5 independent biological repeats.

Our TurboID analysis indicated that SPRY2 interacts with K-RasG12C in a G12Ci-sensitive manner, just as much as well-established effectors C-Raf (Raf1) and B-Raf (**Fig. 1C**). We therefore next verified this again using BRET. In agreement with the TurboID data, the maximal interaction BRET-signal of nL-K-RasG12C and mNG-SPRY2 was dose-dependently reduced after treatment with G12Ci AMG-510/ sotorasib (**Fig. 4E**). Likewise, BRET of K-RasG12V and SPRY2 was dose-dependently reduced after treatment with pan-Ras inhibitor BI-2865 (Kim *et al*, 2023) (**Fig. 4F**).

These data suggest that SPRY2 is recruited to farnesylated, active K-Ras at the plasma membrane via its C-terminal domain, which anchors it there via its palmitoyl moieties.

### SPRY2 binding to K-RasG12V is sensitive to predicted interface mutations

To identify the interface that mediates the potentially direct interaction between K-Ras and SPRY2, we predicted the structure of the complex using AlphaFold 3 (AF3) (Abramson *et al*, 2024). The predicted K-Ras/ SPRY2 complex suggested binding of N-terminal but more so of C-terminal fragment residues of SPRY2 to the effector binding region of K-Ras (**Fig. 5A**). Major putative side-chain contacts between C-terminal SPRY2 residues and K-Ras residues were T298-Q25, K187-Y40 and Y191-R41, respectively (**Fig. 5A**). While these residues were phylogenetically conserved in mouse, chick and mostly also in frog both on the SPRY2 and K-Ras side (**Fig. S6A-C**), conservation amongst other SPRY-family members was limited (**Fig. S6D**), suggesting a specialised role of SPRY2 in this context.

**Figure 5.**
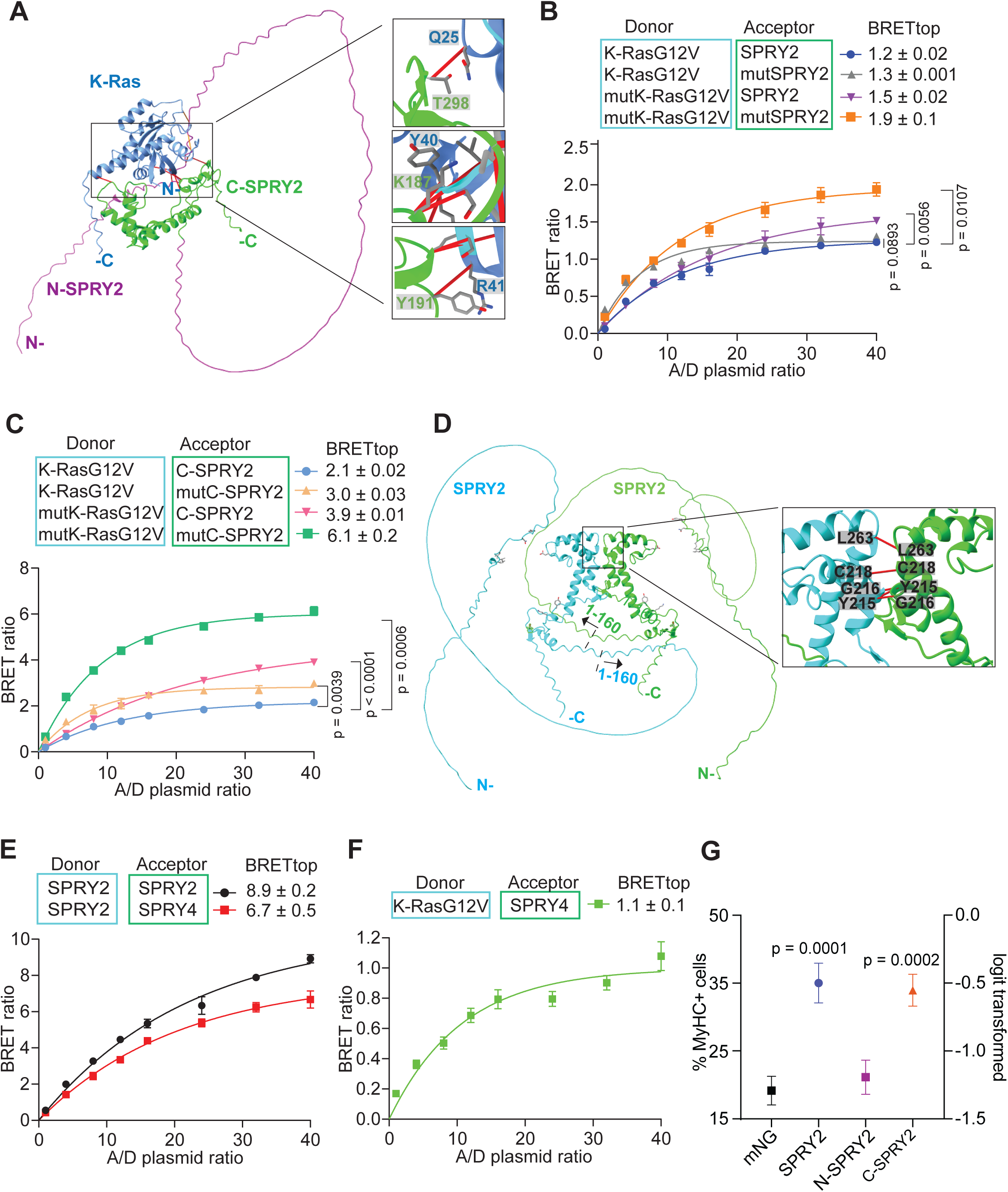
SPRY2 is recruited to active K-Ras on the plasma membrane. (**A**) AlphaFold 3 prediction of K-Ras/ SPRY2 complex, with zoom-in showing residues with putative side chain interactions. (**B,C**) BRET-titration curves of the nanoLuc-tagged K-RasG12V or K-RasG12V-Q25A/Y40A/R41A (mutK-RasG12V) interactions with mNG-tagged SPRY2 or SPRY2-K187A/Y191A/T298A (mutSPRY2) (B), respectively corresponding C-SPRY2 variants (C) in HEK cells from N = 2 to 4 independent biological repeats. (**D**) AlphaFold 3 prediction of the SPRY2/ SPRY2 interaction, with zoom-in showing residues within 3 Å distance. (**E**) BRET-titration curves of the nanoLuc-SPRY2/ mNG-SPRY2 and nanoLuc-SPRY2/ mNG-SPRY4 interaction in HEK cells from N = 4 independent biological repeats. (**F**) BRET-titration curve of the nanoLuc-K-RasG12V/ mNG-SPRY4 interaction in HEK cells from N=3 independent biological repeats. (**G**) Flow cytometric quantification of MyHC terminal differentiation marker expression in C2C12 cells in low serum for 72 hours after transfection with mNG-SPRY2, mNG-N-SPRY2 or mNG-C-SPRY2 constructs. Means ± SD are plotted with N = 3 biological repeats. Statistical analysis was done using one-way ANOVA and Dunn’s post hoc test.

We mutated the six potentially contacting residues in both proteins to alanines and monitored how interaction BRET between them was altered. Surprisingly, interaction of K-RasG12V-Q25A/Y40A/R41A (mutK-RasG12V) with SPRY2-K187A/Y191A/T298A (mutSPRY2) showed a significantly increased BRET (**Fig. 5B**). Also, mixed BRET-pairs of K-RasG12V and mutSPRY2 or mutK-RasG12V and wt SPRY2 showed an elevated BRET-signal (**Fig. 5B**). These differences were even more pronounced when examining these mutations only in the context of C-SPRY2 (**Fig. 5C**). These observations were somewhat explained when examining the AF3-prediction of the mutant complex, which showed an increase in the number of < 3Å contacts to 25, as compared to 13 in the wt complex. These data tentatively support a direct interaction of SPRY2 with the effector binding region of active K-Ras.

The paralog most related to SPRY2 is SPRY1, followed by SPRY4 and SPRY3, however, the expression of the latter is restricted to the brain (Puranik *et al*., 2024). Several other studies of the proximal proteome identified in particular SPRY2 and SPRY4, but not SPRY3, as potential interactors notably of activated K-Ras as compared to the other Ras isoforms (Adhikari & Counter, 2018; Beganton *et al*., 2020; Go *et al*., 2021; Kovalski *et al*., 2019; Song *et al*, 2025). Additionally, SPRY-proteins were suggested to homo-and hetero-di/ oligomerise via their CRD (Ozaki *et al*, 2005). Consistently, AF3-predicted dimers of SPRY2 with a dimer interface that was compatible with concurrent K-Ras binding (**Fig. 5D**). This was supported by BRET-experiments showing a high BRETtop value for the SPRY2/ SPRY2 BRET-pair (**Fig. 5E**). Likewise, BRETtop for the heterodimer of SPRY2/ SPRY4 was high, supporting that SPRY-proteins can hetero-di/oligomerise. In line with a direct or indirect engagement of SPRY4 by active K-RasG12V, the BRET-signal between the two was as high as that with SPRY2 (**Fig. 5F**).

SPRY-proteins have been implicated in various biological contexts as suppressor of Ras-MAPK-signalling. In cancer cells, their role is contentious, with some reporting tumour suppressive activities, while others consider it oncogenic (Masoumi-Moghaddam *et al*, 2014). Current evidence suggests that the normal biological function of SPRY-proteins is associated with stem cell quiescence, such as of muscle cells (Shea *et al*, 2010). However, in the murine C2C12 muscle cell line, SPRY2 expression promotes differentiation (de Alvaro *et al*, 2005). This is in line with the suppression of MAPK-signalling that needs to occur to initiate muscle cell differentiation (Wakioka *et al*, 2001).

We therefore used our well established C2C12 muscle cell differentiation assay to assess the functional effect of mNG-tagged SPRY2 fragment expression (Chippalkatti *et al*, 2024; Parisi *et al*, 2023). Differentiation is quantified by flow-cytometric measurement of the late differentiation marker myosin heavy chain (MyHC). As expected, expression of full length SPRY2 significantly increased differentiation. Consistent with C-SPRY2 binding to and blocking active K-Ras at the plasma membrane (**Fig. 4**), this fragment was sufficient to significantly inhibit differentiation (**Fig. 6G**). By contrast, N-SPRY2, which alone only poorly engages with K-RasG12V (**Fig. 3F**), had no effect on differentiation (**Fig. 6G**).

Based on our data, we propose a model wherein a dimer of SPRY2 with other SPRY proteins is recruited to two active K-Ras proteins by directly binding to their effector binding regions. We further speculate that membrane contacts of the SPRY2/ K-Ras complex localise it in lipid-nanodomains that are also employed by Raf proteins. Thus, SPRY proteins block access of Raf and downregulate specifically Ras-MAPK signalling, which in the muscle C2C12 cell line supports differentiation.

## Discussion

Using proteomics-and BRET-screens, we have identified and categorised two proteins of the proximal K-Ras proteome, which modulate its membrane organisation. Proximity proteomics approaches are inherently noisy. We identified 190 proteins proximal to full length K-Ras proteins. 70 % of these proteins have been identified in at least one previous K-Ras BioID study. However, only six (ARAF, BRAF, CRAF, YKT6, MPP7 and EPHA2) of our proteomics hits were classified as proximal to K-Ras in all previous studies (**Data S1**). Comparison of full-length K-Ras with the minimal K-Ras HVR fragment, tK, that is unable to engage with effectors, together with determination of the sensitivity of the K-RasG12C proximal proteome to G12Ci variously allowed prioritisation of hits for further study. Secondary screening using BRET allowed further stratification of hits. We noticed that the dynamic range in the FLIM-FRET-based hit validation experiments is far greater than in the BRET-screen. However, plate-reader based BRET-screens are easier implemented than microscopy based FLIM-FRET screens (Guzman *et al*, 2016).

We focused our more detailed validation on two hits, APLP2, which is a novel, indirect modulator of K-Ras membrane organisation and SPRY2, which appears to be a direct, effector-like interaction partner of K-Ras. APLP2 engages with C-Raf and may thus impact on the nanoscale organisation of K-Ras, by pre-organising Raf-dimers, which act as Ras nanocluster scaffold (Blazevits *et al*., 2016; Steffen *et al*., 2024).

SPRY-proteins have been implicated as negative regulators specifically of MAPK-signalling (Guy *et al*., 2009). While they are dysregulated in cancer (Masoumi-Moghaddam *et al*., 2014), their normal biological role appears to be associated with the quiescence of stem cells where they suppress MAPK-activity (Barruet *et al*, 2020; Jung *et al*, 2012; Phoenix & Temple, 2010; Shea *et al*., 2010). For instance, re-entry of muscle stem cells into the quiescent state depends on SPRY1-expression (Shea *et al*., 2010). The importance of this process is underscored by the fact that methylation of the SPRY1-gene is observed in muscle cells with aging, which would disrupt re-quiescence of stem cells and consequently foster their demise (Bigot *et al*, 2015).

The C2C12 cell line contains only ∼ 2 % muscle stem cells, hence the quiescent state is less relevant in our assay, where we instead modulate the fate of the committed myoblasts that make up the majority of cells in this cell line (Chippalkatti *et al*., 2024). This is supported by data showing that ablation of SPRY1 decreases the expression of the muscle differentiation marker MyHC in C2C12 cells (Lu *et al*, 2015). Our data are therefore consistent with the SPRY2-mediated downregulation of MAPK-signalling to ectopically initiate cell differentiation (Wakioka *et al*., 2001).

Our mechanistic investigations further suggest that SPRY2 resides in an autoinhibited state in the cytoplasm, which is relieved by the binding of its C-terminal half to active Ras at the plasma membrane.

In line with previous reports, plasma membrane anchorage of SPRY is sensitive to inhibition of palmitoylation, consistent with several C-terminal cysteines that can become palmitoylated (Lim *et al*., 2002; Locatelli *et al*., 2020). We propose that residues of this C-terminal part predominate in making contacts with the effector binding region of K-Ras, however, also N-terminal residues become involved. Indeed, a suppression of MAPK-activity by the C-terminal residues 176-315 of SPRY2 have been reported previously (Egan *et al*, 2002).

The association of SPRY2 with C-Raf seems less prominent but agrees with published results (Sasaki *et al*, 2003). This interaction may occur in different complexes outside of the plasma membrane. Furthermore, we provide evidence that SPRY2 can homo-di-/oligomerise and hetero-di-/oligomerise with SPRY4, consistent with previous reports of their interaction (Rolland *et al*, 2014; Sasaki *et al*., 2003). Given that phylogenetically conserved, predicted interface residues K187, Y191 and T298 of SPRY2 are only moderately conserved in other SPRY-proteins, we propose that ubiquitously expressed SPRY2 serves as the main recruiter of various combinations of SPRY2 homo-and heterodimers. In our proposed model homo-and hetero-di-/oligomers of SPRY2 bind to the effector binding region of active Ras, which blocks access of effectors that prefer the same lipid domains. Thus, Ras activity is essentially clamped down by SPRY-proteins. In this context we suggest that an amphipathic helix at the C-terminus of SPRY2 is positioned to insert into the membrane and mediates the lipid selectivity. The complex of dimeric Ras that is scaffolded on SPRY-dimers would explain the observed BRET-signature that was indicative of SPRY-dimers stabilizing dimeric Ras nanocluster. Currently, only the structure of short phosphorylated SPRY2 and SPRY4 peptides centred around Y55 bound to a c-Cbl TKB domain exists (Ng *et al*, 2008; Sun *et al*, 2010).

Our study underscores the need for comprehensive structural studies of SPRY/ Ras complexes to examine our structural predictions experimentally and for studies elaborating the exact mechanism and biological context of SPRY-mediated Ras-MAPK-pathway suppression.

## Materials and Methods

All equipment, software, materials and reagents and their sources are listed in **Table S1**.

### Expression Constructs

TurboID constructs were generated by Genscript in pcDNA3 vector under the control of either CMV or PGK promoter. The design included an N-Terminal HA-tag, followed by the linker GSGSGGSG, then the TurboID sequence separated from the K-Ras sequence by linker GGSGSGSG (Cho *et al*, 2020).

Multi-site Gateway cloning was used to generate plasmids encoding mNeonGreen-tagged (mNG) K-RasG12V and K-RasG12C as well as the control vector containing only mNG, as described in the previous studies by us (Manoharan *et al*., 2023; Okutachi *et al*., 2021; Wall *et al*, 2014). The entry clone plasmid encoding mNG was synthetised by GeneCust. In brief, three entry clones, encoding the CMV promoter, the mNG tag and the gene of interest or a stuffer sequence for the control vector, with compatible LR recombination sites, were recombined with a destination vector, pDest-305 or pDest-312, using the Gateway LR Clonase II enzyme mix (Thermo Fisher Scientific, #11791020). The Gateway reaction mix was then transformed into the ccdB-sensitive *Escherichia coli* strain DH10B (Thermo Fisher Scientific, #EC0113), and positive clones were selected using ampicillin. All final clones were verified using Sanger sequencing by Eurofins.

### Cell culture

HEK293-EBNA cells were cultured in Dulbecco’s modified Eagle Medium (DMEM) (ThermoFisher Scientific, #41965039) supplemented with ∼ 9 % (v/v) Fetal Bovine Serum (FBS) (ThermoFisher Scientific, A5256701), 2 mM L-Glutamine (ThermoFisher Scientific, #25030024) and penicillin-streptomycin (ThermoFisher Scientific, #15140122) 10,000 units/ mL, in T75 (Greiner, #658175) or T175 (Greiner, #660175) flasks. Cells were cultured in a humidified incubator maintained at 37°C and 5 % CO_2_ and routinely passaged 2-3 times per week.

HEK293T were a gift from Horizon Discovery, HEK293 were purchased from ATCC (CRL-1573). HEK293-TurboID stable cell lines were generated by transfection and selection of stable transformants with G418. Single clones were not isolated. For SILAC labelling, cells were grown for five passages in DMEM supplemented with 10 % dialysed FBS (BioSera, #FB-1001D) containing light, medium or heavy-labelled arginine and lysine (Cambridge Isoptope Laboratories). Labelling was confirmed to be ≥ 95 % using liquid chromatography mass spectrometry (LC-MS) prior to experiments.

C2C12 cells were cultured in DMEM supplemented with ∼ 9 % (v/v) FBS, 2 mM L-and penicillin-streptomycin 10,000 units/ mL (high serum medium). Cells were cultured in a humidified incubator maintained at 37°C and 5 % CO_2_ and passaged at about 50 % confluency. To induce differentiation, the medium was exchanged with DMEM supplemented with ∼ 2 % horse serum (ThermoFisher Scientific, #16050130), 2 mM L-glutamine and penicillin/ streptomycin at 10,000 units/ mL (low serum medium).

### Western blotting

For SDS-PAGE analysis of TurboID experiments, NuPage 4-12 % Bis-Tris gels (ThermoFisher, #NP0321, #NP0336, #WG1402) and MOPS running buffer (ThermoFisher, #NP0001) were used. The gels were transferred to a nitrocellulose membrane (Amersham Protran 0.45 µm, #10600002) using Genie Blotter (Idea Scientific, #4017) and the membranes were blocked either in 5 % milk powder (Marvel) or 5 % bovine serum albumin (BSA) both diluted in Tris-buffered saline (TBS) supplemented with 0.1 % TWEEN 20. Primary antibodies were diluted in a blocking buffer and incubated at 4 °C overnight, secondary antibodies were diluted 1:15000 in TBS with 0.1 % TWEEN 20 and 5 % milk and incubated for 1 hour at room temperature (20-25°C). Streptavidin-800CW conjugate was diluted at 1:5000 and incubated as per secondary antibodies. Amersham ECL Full-Range Rainbow Marker (Cytiva, #RPN800E) was used as a protein size reference.

For the remaining experiments, Mini-PROTEAN TGX TM Precast Protein Gels 4-20 % (BioRad, #4561094 and #4561093) were used. Page Ruler Prestained Protein Ladder (Thermo Scientific, #26616) was used as a protein size reference. For analysis by Western blot, the samples were transferred to a 0.2 µm nitrocellulose membrane using Trans-Blot Turbo RTA Midi-size Nitrocellulose Transfer Kit (BioRad, #1704271). The membranes were rinsed with water and then blocked in TBS supplemented with 0.2 % Tween 20 and 2 % BSA (ITW Reagents, #A6588, 0100) for 1 h at room temperature (20-25°C). Primary antibodies were diluted in a blocking buffer and were incubated overnight at 4°C. All secondary antibodies were diluted 1:10,000 in a blocking buffer and were incubated 1 h at room temperature (20-25°C).

Membranes were visualised using an Odyssey CLx Imaging System. Quantitation of bands was performed using ImageStudioLite software (v5.2.5) and graphs were plotted using Graph Pad Prism 10.4.2.

### TurboID experiments

For the first experiment, cells were grown in SILAC media for mass spectrometry experiments (“Light”= unlabelled; “medium” = Lysine 4 [4,4,5,5 – D_4_], Arginine 6 [^13^C_6_]; “heavy” = Lysine 8 [^13^C_6_, ^15^N_2_], Arginine 10 [^13^C_6_, ^15^N_4_]). HEK293T cells were transfected using the calcium phosphate method. Media was changed 1 h prior to transfection. For a 15 cm dish, 17.5 μg DNA was mixed with 75 μL 2.5M CaCl_2_ and diluted to 700 μL using 0.1x TE (1 mM Tris-HCl pH7.6, 0.1 mM EDTA) before adding dropwise to 700 μL 2 x HBS (50 mM HEPES pH7.05, 140 mM NaCl, 1.5 mM Na_2_PO_4_ while vortexing. Complexes were allowed to form for 3 min at 37 °C with shaking at 600 rpm, before adding dropwise to cells. After 3 h incubation at 37 °C, cells were washed and media was changed, and cells were incubated for approximately 24 h before treatment for experiments. TurboID pull-downs were performed as described elsewhere (Cho *et al*., 2020). Transfected cells were treated with 50 μM biotin (Sigma-Aldrich, #B4501) or DMSO for 10 min at 37 °C where indicated, before washing three times in ice cold PBS and lysis in RIPA buffer (50 mM Tris pH 7.5, 150 mM NaCl, 0.1 % SDS, 0.5 % sodium deoxycholate, 1 % Triton X-100) supplemented with 50 mM NaF, PhosSTOP (Roche, #04 906 837 001), 2 mM sodium orthovanadate, 0.1 mM phenylmethylsulfonyl fluoride (PMSF) and mammalian protease inhibitors (Sigma-Aldrich, #P8340). Lysates were sonicated, clarified by centrifugation and the concentration determined by Pierce BCA protein assay (Thermo Scientific, #A55864). Lysates were mixed 1:1:1 (light:medium:heavy) and 3 mg pulled down using 250 μL streptavidin magnetic beads (Thermo Scientific, #88817). Beads were washed with 2 × RIPA, 1 × 1 M KCl, 1 × 0.1 M Na_2_CO_3_, 2 M Urea/ 10 mM Tris-HCl pH 8.0, 2 × RIPA, 1 × 50 mM Tris pH 7.5, 2 × 2 M Urea/ 50 mM Tris pH 7.5. For digestion, beads were incubated with 400 ng Trypsin Gold (Promega, # V5280) in 2 M urea/Tris pH 7.5 for 1 h at 25 °C. The digest solution was removed from the beads and the digestion allowed to continue overnight before reduction, alkylation and dehydration.

For the second experiment, SILAC-labelled HEK293 cells stably expressing TurboID constructs were treated with K-RasG12C inhibitors or vehicle for 3 h and 50 min at 37 °C where indicated, prior to 10 min of biotin treatment. Inhibitor concentrations were 1 μM AMG510, 10 μM ARS1620 or 3 μM MRTX849. Lysis was as above, then a total of 900 μg lysate was pulled down with 75 μL strepatavidin beads as before. Beads were boiled at 95°C in 3 × protein loading buffer (165 mM Tris pH 6.8, 17 % glycerol, 5 % SDS, 0.045 % DTT, 0.6 % bromophenol blue) supplemented with 2 mM biotin and 20 mM freshly added DTT, then loaded onto an SDS-PAGE gel. Peptides were yielded by in-gel trypsin digest. All LC-MS samples were desalted using C18 spin columns prior to LC-MS analysis (Thermo Scientific, #89870).

TurboID pulldowns for Western blotting were performed as per experiments 1 or 2, but without SILAC labelling.

### LC-MS analysis

In experiment 1, the Ras proximal proteins were analysed by Warwick Scientific Service, University of Warwick. Tryptic peptides from three biological replicates were analysed using an UltiMate 3000 RSLCnano system coupled to a Thermo Orbitrap Fusion (Q-OT-qIT, Thermo Scientific). Peptides were loaded onto a trapping column (Acclaim PepMap µ-precolumn, 300 µm × 5 mm, 5 μm packing material, 100 Å; ThermoFisher Scientific) in 4 % acetonitrile/ 0.1 % formic acid, then eluted onto the analytical column (Acclaim PepMap RSLC 75 µm × 50 cm ^2^ µm 100 Å; ThermoFisher Scientific) by increasing from 4 % acetonitrile/ 0.1 % formic acid to 25 % acetonitrile/ 0.1 % formic acid over 36 min, then to 35 % acetonitrile/ 0.1 % formic acid over 10 min, and to 90 % acetonitrile/ 0.1 % formic acid over 3 min, followed by a 10 min re-equilibration at 4 % acetonitrile/ 0.1 % formic acid. Eluting peptides were ionised using an electrospray source and analysed on a Thermo Orbitrap Fusion (Q-OT-qIT, Thermo Scientific). MS survey scans from 375 to 1575 *m*/*z* were acquired at 120K resolution (at 200 *m*/*z*) with a 50 % normalised automatic gain control (AGC) target and a max injection time of 150 ms. Precursor ions with charge state of 2-6 were selected for MS/MS by isolation at 1.2 Th. Higher-energy collisional dissociation (HCD) fragmentation was performed with normalised collision energy of 33, and rapid scan MS analysis in the ion trap. The MS^2^ was set to 50 % normalised AGC target and the maximum injection time was 150 ms. The dynamic exclusion duration was set to 45 s with a 10 ppm tolerance. Monoisotopic precursor selection was turned on. The instrument was run in top speed mode with 2 s cycles.

In experiment 2, the G12Ci sensitive proteome was analysed in the Centre for Proteomics Research, University of Liverpool. Peptides from three biological replicates were analysed using an UltiMate^™^ 3000 RSLCnano system coupled to a Q Exactive^™^ HF Hybrid Quadrupole-Orbitrap^™^ Mass Spectrometer (ThermoFisher Scientific). Peptides were loaded onto a trapping column (Acclaim PepMap 100 C18, 75 μm × 2 cm, 3 μm packing material, 100 Å; ThermoFisher Scientific) using 0.1 % trifluoroacetic acid, 2 % acetonitrile in water at a flow rate of 12 μL min^−1^ for 7 min. Peptides were eluted onto an analytical column (EASY-Spray PepMap RSLC C18, 75 μm × 50 cm, 2 μm packing material, 100 Å) at 30 °C using a linear gradient of 90 min rising from 3 % acetonitrile/ 0.1 % formic acid to 40 % acetonitrile/ 0.1 % formic acid at a flow rate of 300 nL min^−1^. The column was then washed with 79 % acetonitrile/ 0.1 % formic acid for 5 min, and re-equilibrated to starting conditions. The nano-liquid chromatograph was operated under the control of Dionex Chromatography MS Link 2.14. The nano-electrospray ionisation source was operated in positive polarity under the control of QExactive HF Tune, with a spray voltage of 1.8 kV and a capillary temperature of 250 °C. Full MS survey scans between m/z 350-2,000 were acquired at a mass resolution of 60,000 (full width at half maximum at m/z 200) with AGC target set to 3e^6^, and a maximum injection time of 100 ms. The 16 most intense precursor ions with charge states of 2– 5 were selected for MS^2^ with an isolation window of 2 m/z units. HCD was performed to fragment the selected precursor ions using a normalised collision energy of 30 %. Product ion spectra were recorded between m/z 200-2,000 at a mass resolution of 30,000 (full width at half maximum at m/z 200). For MS^2^, the AGC target was set to 1e^5^, and the maximum injection time was 45 ms. Dynamic exclusion was set to 30 s.

### LC-MS Data analysis

The raw data were searched using MaxQuant version 1.6.17 against a human Swissprot and tREMBL database (experiment 1) or version 2.2.0 against a human Swissprot database (experiment 2). For the database search, peptides were generated from a tryptic digestion with up to two missed cleavages, carbamidomethylation of cysteines as fixed modification. Oxidation of methionine, biotinylation of lysine and acetylation of the protein N-terminus were added as variable modifications. Further analysis of the data was performed using Excel and Perseus (v1.6.15.0 for experiment 1, v2.0.9.0 for experiment 2). Protein IDs that matched the reverse or known contaminants list were excluded. To allow for ratio generation in Excel, a nominal value of 1,000 was added to all intensities. Protein IDs that were present in at least two repeats for one construct were retained. To refine a shortlist for continuation to BRET-assays, mean ratios (+biotin/-biotin) of intensities for protein IDs from all 3 repeats of experiment 1 were generated and only hits increased by at least 2-fold compared to the -biotin condition in full-length TurboID-KRAS constructs but not TurboID-tK or TurboID alone were retained. This list was combined with common hits from previously published Ras BioID experiments and controls to generate a shortlist of proteins of interest for inclusion in BRET-based proximity assays (Adhikari & Counter, 2018; Beganton *et al*., 2020; Kovalski *et al*., 2019).

For experiment 2, a proximity list was defined as any proteins enriched at least 2-fold in the TurboID-G12C plus biotin condition vs TurboID-G12C minus biotin. Of this group, G12Ci-sensitive hits were defined as any that were less enriched in TurboID-G12C plus G12Ci plus biotin, vs TurboID-G12C plus biotin. Only hits with at least two values returned for at least two of any of the G12Ci conditions were retained.

### GFP-Trap mediated pull-downs

HEK293-EBNA cells were plated in a 10 cm dish scale so that for transfection they would be 50-70 % confluent (2 × 10^6^ cells per dish for next day transfection). For each sample, one 10 cm dish was plated. For transfection, 5 µg of each DNA to be transfected was diluted in 500 µL jetPRIME buffer. For single-plasmid transfections, jetPRIME was used at 3:1 ratio (3 µL of jetPRIME per 1 µg of DNA transfected) and for plasmid co-transfection at 2:1 ratio (2 µL of jetPRIME per 1 µg of DNA). After approximately 24 h, the growth medium was removed and the cells were rinsed gently twice with cold PBS (DPBS, Gibco, #14040091). Next, 500 µL of the lysis buffer (2 mM EDTA, 50 mM Tris pH 7.5, 150 mM NaCl, 0.5 % NP-40, Pierce Protease Inhibitor Tablet EDTA-free (Thermo Scientific, #A32955), PhosSTOP (Roche, #04 906 837 001, and 1 mM DTT) was applied to each dish, the cells were scraped, transferred to an Eppendorf tube and incubated on ice for 30 min. The samples were mixed by gently inverting 3-4 times. The lysate was cleared by centrifuging for 15 min at 4°C and ∼ 16,500 × *g*. Cleared lysate was transferred to a clean Eppendorf tube and subjected to GFP-Trap pull-down (GFP-Trap Agarose, ChromoTek #gta-20). For pull-downs, 25 µL bead slurry per 10 cm cell dish were considered. The required volume of bead slurry was pipetted to an Eppendorf tube and rinsed three times with equilibrating buffer (10 mM Tris pH 7.5, 150 mM NaCl, 0.5 mM EDTA). The beads were finally re-suspended in the equilibrating buffer to have 1:1 bead:buffer ratio. Pull-downs were conducted for 3 h at room temperature (20-25°C) with samples in gentle rotation. Next, the beads were pelleted by centrifuging at 4°C and 800 2 × *g* for 1 min, supernatant was removed and the beads were rinsed with 1 mL washing buffer (2 mM EDTA, 50 mM Tris pH 7.5, 150 mM NaCl, 0.5 % NP-40, 1 mM DTT), at 4°C for 15 min on a rotating wheel. The washing step was repeated for total of four times. The bound material was released from the beads by adding 25 µL of 2 × SDS-PAGE sample buffer and incubating at 95°C for 5 min. Supernatant was recovered by centrifuging at 800 × *g* for 1 min and transferred to a clean Eppendorf tube. The samples were resolved on 4-20 % SDS-PAGE gels in Tris-Glycine buffer and further analysed by Western blotting. For each sample, 2 % of total lysate was loaded as input control.

### Bioluminescence resonance energy transfer (BRET) assays

For screening the effect of TurboID hits siRNA-mediated knockdown in HEK293-EBNA cells, we employed BRET-assays with RLuc8-fused BRET donor, GFP2-fused BRET acceptor and coelenterazine 400a as the substrate (Duval *et al*., 2024). A CLARIOstar plate reader (BMG Labtech) was used for measurements. The cells were plated in a 12-well cell culture plate (Greiner Bio-One, #655180). The following day, the cells were transfected with siRNA at the final concentration of 100 nM using Lipofectamine RNAiMAX diluted in Opti-MEM (ThermoFisher Scientific, #31985047), while the cells were left in complete DMEM. After ∼ 24 h the siRNA-containing medium was replaced with fresh DMEM and ∼ 1 µg BRET-sensor constructs was transfected per well at the donor:acceptor plasmid ratio of 1:15 (65 ng of the BRET-donor plasmid) and using 3 µL jetPRIME transfection reagent. The pcDNA3.1(-) control plasmid was used to top up the DNA amount.

The following day, the control samples were treated with 10 µM mevastatin, or DMSO as vehicle control at 0.1 % (v/v) for 16-18 h. About 48 h after transfection of BRET-sensors, the cells were plated in a white flat bottom 96-well plate (Thermo Fisher Scientific, #236108) and BRET measurements were taken.

First, the fluorescence intensity of GFP2 was measured (*λ*_excitation_ 405 ± 10 nm and *λ*_emission_ 515 ± 10 nm). This signal is directly proportional to the expression of the BRET-acceptor (RFU). Next, coelenterazine 400a was prepared by diluting in PBS to 100 µM concentration, and 10 µL was dispensed to each well, thus making its final concentration in the well 10 µM. BRET was recorded simultaneously at *λ*_emission_ 410 ± 40 nm (RLU) and 515 ± 15 nm (BRET signal). RLU is directly proportional to the expression of the BRET-donor. BRET ratio was calculated as the ratio of BRET signal/ RLU and the final BRET ratio was obtained by subtracting the background BRET ratio obtained from cells expressing only BRET-donor plasmid:

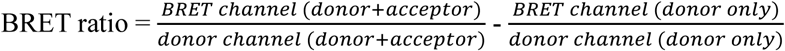

The percentage modulation on BRET ( ΔBRET) is calculated as:

ΔBRET=|100−[100∗(Mean BRET ratio of sample/Mean BRET ratio of scrRNA)]|. A negative sign is added if the gene decreases the BRET ratio. Heatmaps were generated in Prism from calculated averaged BRET ratios across all biological repeats.

Interaction of SPRY2 fragments with K-RasG12V, mutK-RasG12V, K-RasG12C, H-RasG12V, SPRY2 and SPRY4 was assessed by BRET titration experiments using a modified pair of BRET-sensors, with nanoLuc (nLuc)-fused BRET-donor and mNeonGreen (mNG)-fused BRET acceptor. The concentration of BRET-donor plasmid was kept constant (25 ng) and BRET-acceptor plasmid was titrated to 1:40 donor:acceptor plasmid ratio. The pcDNA3.1(-) plasmid was used to top-up plasmid amount in each well to ∼ 1 µg and 3 µL of jetPRIME transfection reagent were used per well.

Clariostar settings were adjusted as follows: for fluorescence intensity (RFU), *λ*_excitation_ 485 ± 10 nm and *λ*_emission_ 535 ± 10 nm, and for BRET measurements, *λ*_emission_ 460 ± 25 nm (RLU) and 535 ± 25 nm (BRET signal). Coelenterazine a was used at 2.9 µM final concentration. Compounds were prepared at the indicated concentrations by diluting in DMEM and 1 mL of the mix was added to each well, for the total of 18-24 h, with DMSO 0.05 % or 0.01 % (v/v) as vehicle control.

The BRET ratio was plotted against acceptor:donor (A/D) plasmid ratio and was fitted using one-phase association fit in Prism. The BRETtop corresponds to the highest BRET ratio reached at acceptor:donor = 40:1 plasmid ratio and is given as mean with standard error (SEM).

### FLIM-FRET measurements

A total of 80,000 HEK293T cells were seeded in 12-well plates. Each well contained a sterilized 16-mm coverslip and 1 mL of medium. The next day, the cells were transfected using jetPRIME at 1:1 ratio (1 µL of jetPRIME for 1 µg of DNA). For donor-only samples, the cells were transfected with 500 ng of pmEGFP-tagged K-RasG12V plasmid. For K-RasG12V nanoclustering-FRET, the cells were transfected with a total of 1 µg of pmEGFP/pmCherry-K-RasG12V, at a donor: acceptor plasmid ratio of 1:3. For gene silencing experiments, the cells were co-transfected with 100 nM siRNA along with the FRET plasmids using jetPRIME. After four h, the media was changed. After 48 h of transfection, the cells were fixed in 4 % paraformaldehyde for 10-15 min. The cells were then mounted on slides using Mowiol 4-88. The lifetime of the mGFP donor was measured per cell using a fluorescence microscope (Zeiss AXIO Observer D1) with a lifetime imaging attachment from Lambert Instruments. The FRET efficiency (E) was calculated using the formula: E = (1 − τDA / τD) × 100 %, where τDA is the lifetime of a FRET condition averaged across all acquired samples, and τD is the averaged lifetime of the donor-only sample.

### Confocal Microscopy

HEK293-EBNA cells were seeded at 350,000 cells in 2 mL of complete DMEM onto Glass coverslips 1.5H (Carl Roth, Karlsruhe, Germany, #LH22.1) in 6-well plates (Greiner Bio-One, #657160). The following day, the cells were transfected with 0.25 μg of plasmid diluted in 200 μL jetPRIME buffer and 2 μL jetPRIME transfection reagent. For double transfection when the total plasmid amount was 0.5 µg, the same amount of buffer and jetPRIME reagent were used. Compound treatments were done 6 h after transfection by recovering the medium from each well in Eppendorf tubes and diluting the compounds to the indicated concentrations. After 18 h of treatment, the medium was removed and the cells were rinsed with PBS. Next, the cells were fixed using 4 % paraformaldehyde (ThermoFisher Scientific, #43368.9M) in PBS and left to incubate for 10 min at 20-25 °C. The fixing solution was removed and the coverslips were covered with a 1:2,000 dilution of Hoechst dye (ThermoFisher Scientific, #H1399) for 5 min. After removal of Hoechst dye, the cells were washed twice with PBS. The coverslips were then mounted using VECTASHIELD Mounting Medium (Vector Laboratories, #H-1000-10) and left to dry. The slides were observed on a Nikon Eclipse Ti2-E spinning disk confocal microscope with an iXon Ultra 888 EMCCD camera (Andor, Oxford Instruments) using a plan APO 60×/1.40 Ph3 DM oil immersion objective (Nikon, Belgium) and NIS-Elements Imaging Software (Nikon, Version 6.10.01). The mNeonGreen-label was detected using 488 nm excitation and detected using EGFP emission settings (535/20 band pass filter). The mCherry-label was detected using 561 nm excitation and a 560/40 band pass filter for signal detection. Images were analyzed using Fiji ImageJ2 (v2.16.0).

### Flow cytometry-based C2C12 cell differentiation assay

The differentiation assay was performed on a flow cytometer Guava easyCyte HT-2L flow cytometer as described by us previously (Chippalkatti *et al*., 2024; Parisi *et al*., 2023). In brief, C2C12 cells were seeded in 6-well plates at a density of 20,000 cells in 2 mL. The next day, cells were transiently transfected at 60 % confluency using jetPRIME at a 3:1 ratio (3 µL of jetPRIME per 1 µg of DNA) with 2 µg of mNG-SPRY2, mNG-N-SPRY2 or mNG-C-SPRY2 diluted in 200 µL of jetPRIME buffer. The medium was replaced by fresh medium 4 h later. 24 h after transfection, the wells were switched to low serum DMEM containing ∼ 2 % horse serum. The medium was changed every day for three days. Then, cells were harvested by trypsinisation for 5 min and centrifuged at 500 × *g* for 5 min. The cell pellet was fixed with 4 % paraformaldehyde (ThermoFischer Scientific, #43368.9M) in PBS for 10 min. After washing with PBS, the cells were permeabilised with 0.5 % Triton X-100 in PBS for 10 min. Next, the cells were washed with PBS containing 0.05 % Tween-20 (PBST) and immunolabelled with eFluor 660-conjugated anti-myosin 4 (myosin heavy chain, MyHC) antibody (ThermoFisher Scientific, #50-6503-82), at 1:100 dilution in PBST for 1 h at 4 °C. The cells were then centrifuged at 500 × *g* for 5 min and resuspended in PBST for flow cytometric analysis. Non-labelled, mNG-only expressing and MyHC-positive cells were used to establish gates and detection channel settings. The mNG-variants were detected by 488 nm excitation and the Grn-B (525/30) band pass filter. MyHC immunolabelled with eFluor 600-conjugated antibody was detected by 640 nm excitation and the Red-R (664/20) band pass filter. Typically, > 1000 cells were analysed in the final target gate per experimental repeat. Intact cells were quantified for the expression of mNG in the mNG-low bin (up to 10-fold above background fluorescence) and MyHC using the ‘Quad Stat plot’ feature on the GuavaSoft 4.0 software.

### Statistics and Data Analysis

Data were analysed using GraphPad Prism10.4.2 software. The number of independent biological repeats (N) is indicated in the figure legends. Plotted are means with standard error (SEM) unless stated otherwise.

The statistical evaluations of the Western blotting quantifications were calculated using unpaired t-test with Welch’s correction. All BRETtop values were compared using One-Way Brown-Forsythe and Welch ANOVA tests with Dunnett’s T3 correction for multiple comparisons using GraphPad Prism. Other statistical tests are described in the legends. A p-value of < 0.05 was considered statistically significant and p values are annotated in the figures. In addition, statistical significance levels may be annotated in the plots as *: p < 0.05; **: p < 0.01; ***: p < 0.001; ****: p < 0.0001.

## Acknowledgements

LC-MS samples were run by Warwick Scientific Services (University of Warwick, UK) and the Centre for Proteome Research (University of Liverpool, UK). We thank Dr. Cleidi Zampronio and Dr. Philip Brownridge for their assistance with these experiments.

This work was supported by grants from the BBSRC (BB/T012757/1) to IAP and the Luxembourg National Research Fund (FNR) AFR/17927850/Duval C./ Kruptor to CJD and INTER/UKRI/19/14174764 to D.K.A.

## Disclosure statement and competing interests

The authors declare that they have no known competing financial interests or personal relationships that could have appeared to influence the work reported in this paper.

## Supplementary Information

### Supplementary Figure Legends

**Figure S1.**
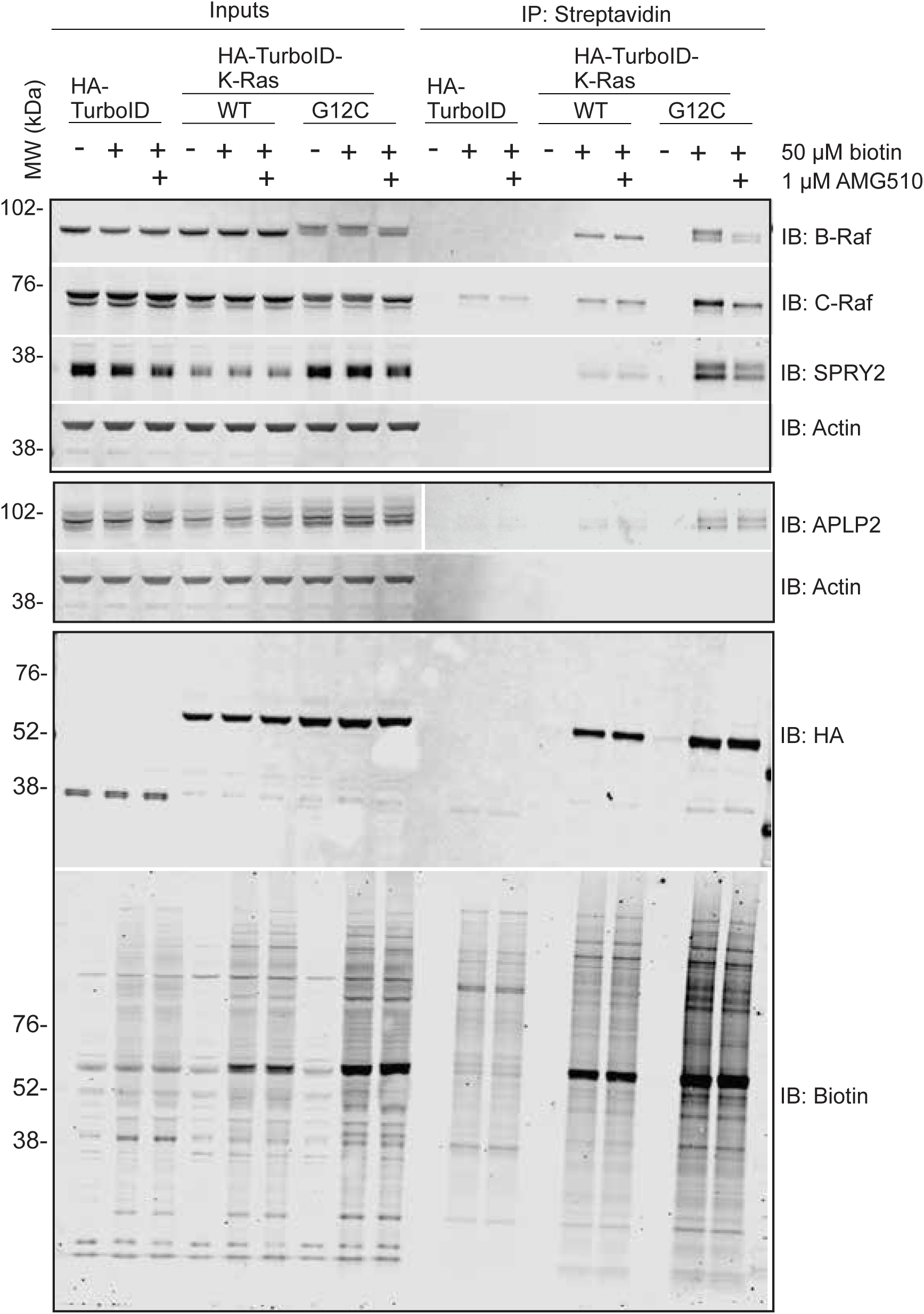
Related to main Figure 1. Representative Western blots of the K-Ras proximal labelling and its activation dependence for a subset of hits showing successful biotinylation, streptavidin enrichment, K-RasG12C inhibitor (G12Ci) efficacy and activation dependence of selected hits. Input versus post-streptavidin enrichment of biotinylated proximal proteins using the same TurboID experiment configuration employed for mass spectrometry analysis (N = 3).

**Figure S2.**
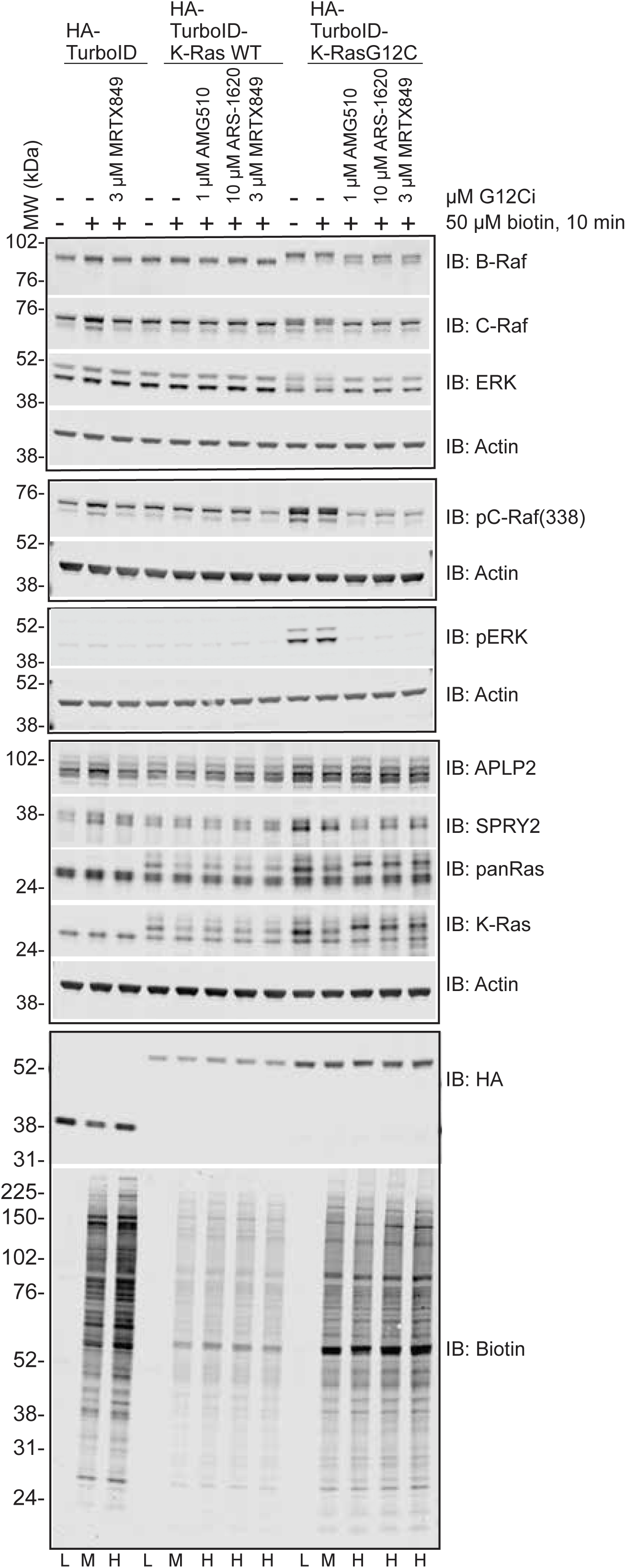
Related to main Figure 1. Activation-dependent changes with a panel of K-Ras inhibitors in stable HEK293-TurboID cell lines. Responses to three different covalent K-RasG12C inhibitors (G12Ci) are shown. The lysates come from one of the total of N=3 repeats of the TurboID streptavidin pull-downs and mass spectrometry analysis of SILAC-labelled cells.

**Figure S3.**
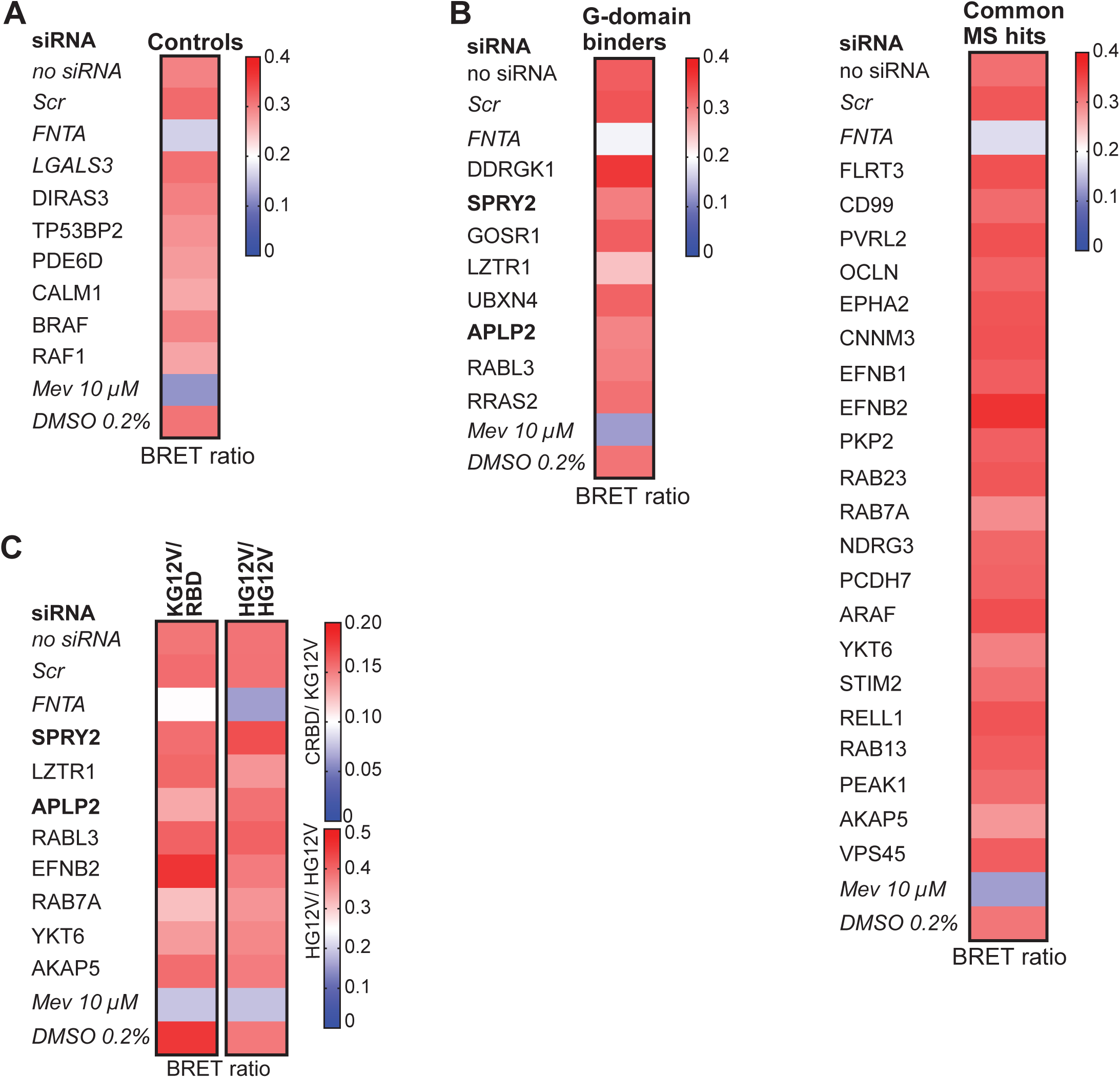
Related to main Figure 1. (**A,B**) Heatmaps of BRET ratio values after knockdown of the indicated genes in the BRET-assay to measure K-RasG12V membrane organisation in HEK cells (donor:acceptor plasmid ratio = 1:15. The control set of genes (A) and our selected TurboID hits combined with hits from our meta-analysis (B) were examined. Data show means from N ≥ 4 independent biological repeats. (**C**) Heatmaps of BRET ratio values after knockdown of the indicated genes in BRET-assays to measure K-RasG12V/ C-Raf-RBD interaction (KG12V/ RBD, donor:acceptor plasmid ratio = 1:15) and H-RasG12V membrane organisation (HG12V/ HG12V, donor:acceptor plasmid ratio = 1:15) in HEK cells. Data show means from N = 4 independent biological repeats.

**Figure S4.**
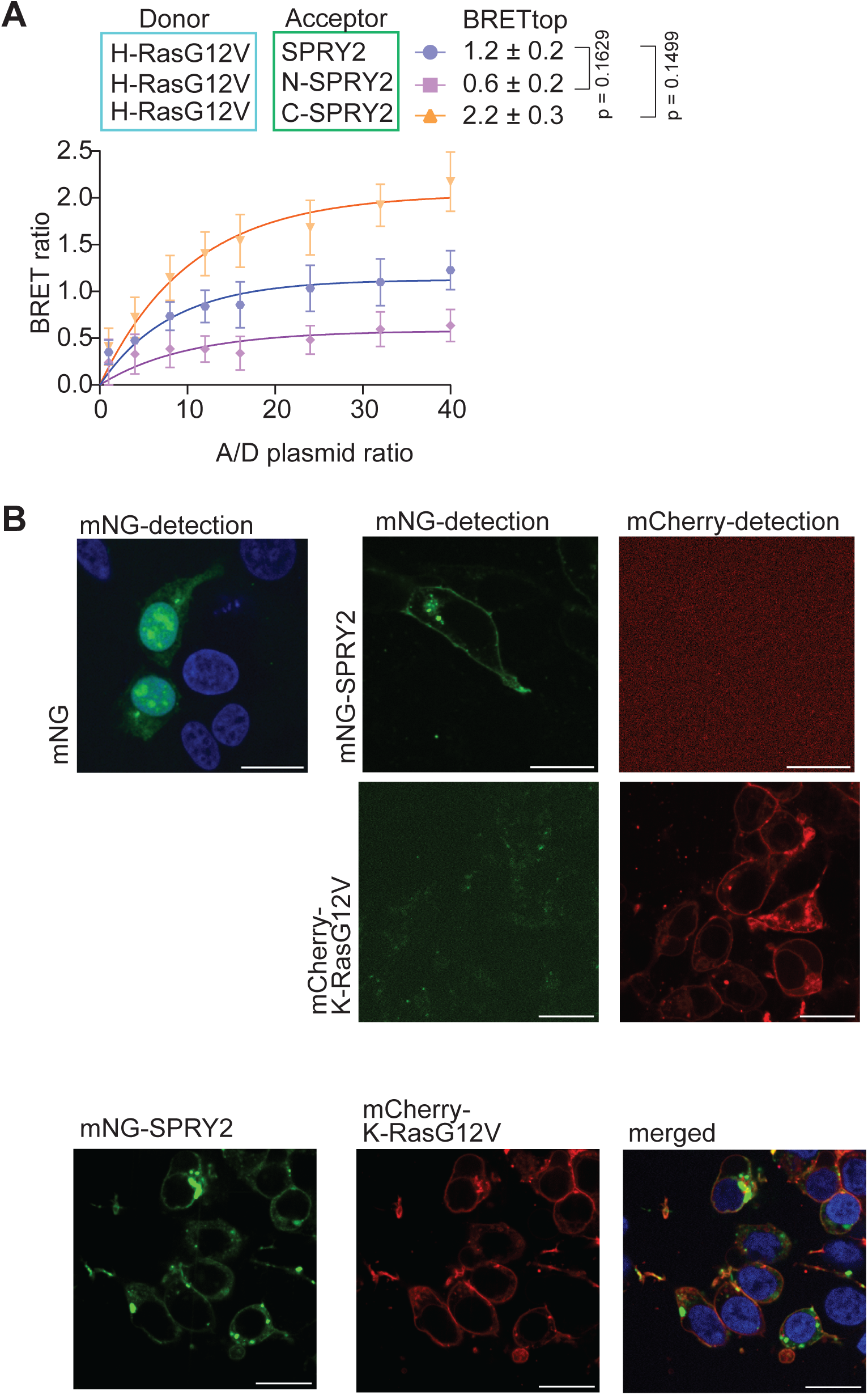
Related to main Figure 3. (A) BRET-titration curves of the nanoLuc-H-RasG12V interaction with mNG-tagged SPRY2 fragments in HEK cells from N = 3 biological repeats. (B) Confocal imaging showing expression of mNG without fused protein for reference (top left), imaging crosstalk controls (top right), and co-localisation of mCherry-K-RasG12V with mNG-SPRY2 in HEK cells (bottom). Scale bar = 20 µm.

**Figure S5.**
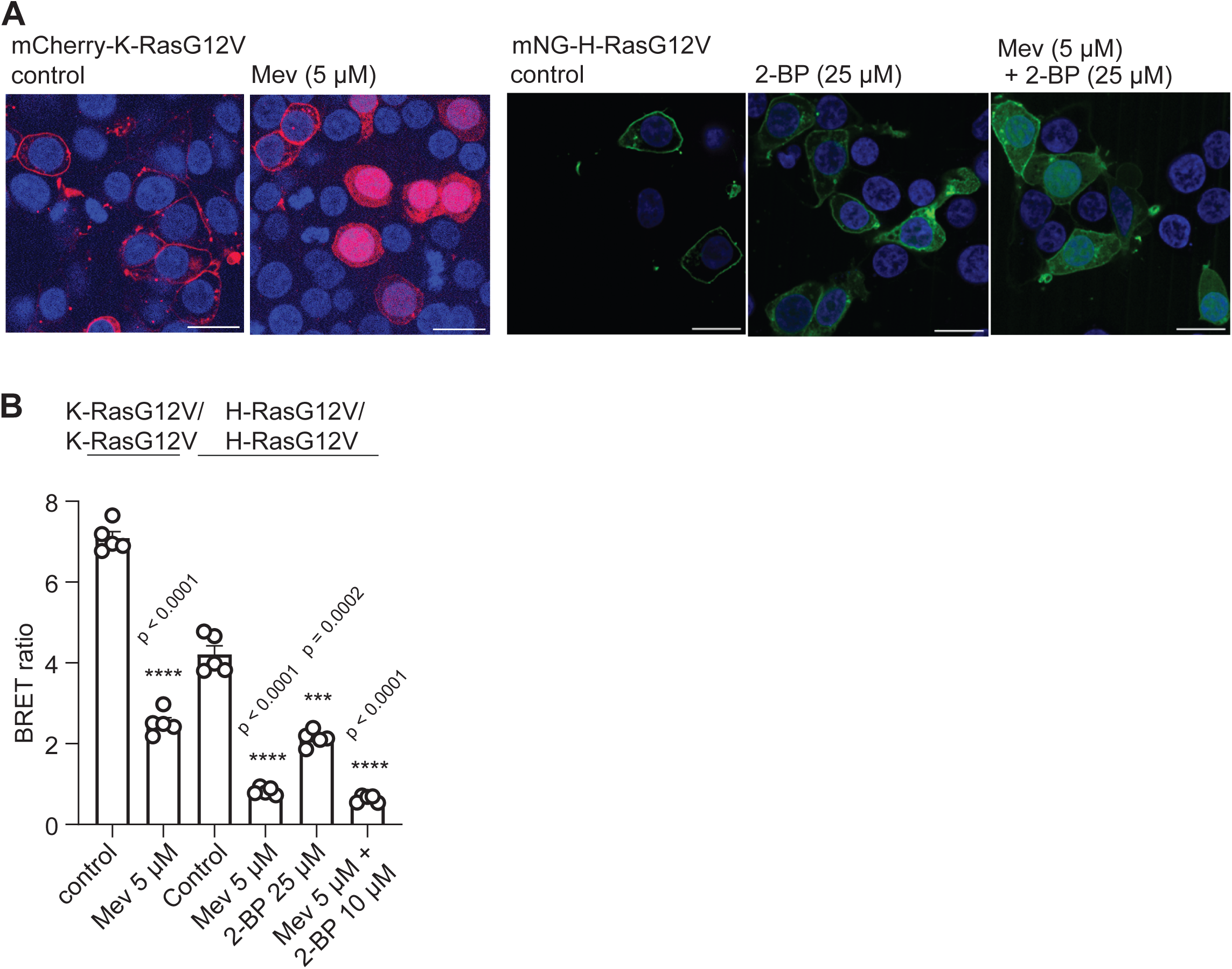
Related to main Figure 4. (A) Confocal imaging showing changes in subcellular distribution of mCherry-K-RasG12V (left) or mNG-H-RasG12V (right) in HEK cells following treatment with indicated lipidation inhibitors mevastatin (Mev) and/ or 2-bromopalmitate (2-BP). Scale bar = 20 µm. (B) Effect of mevastatin (Mev) and/ or 2-bromopalmitate (2-BP) on interaction BRET of nano-luc-K-RasG12V/ mNG-K-RasG12V (donor:acceptor plasmid ratio = 1:8) and nanoLuc-H-Ras-G12V/ mNeonGreen-H-RasG12V (donor:acceptor plasmid ratio = 1:5) in HEK cells. N = 5 independent biological repeats.

**Figure S6.**
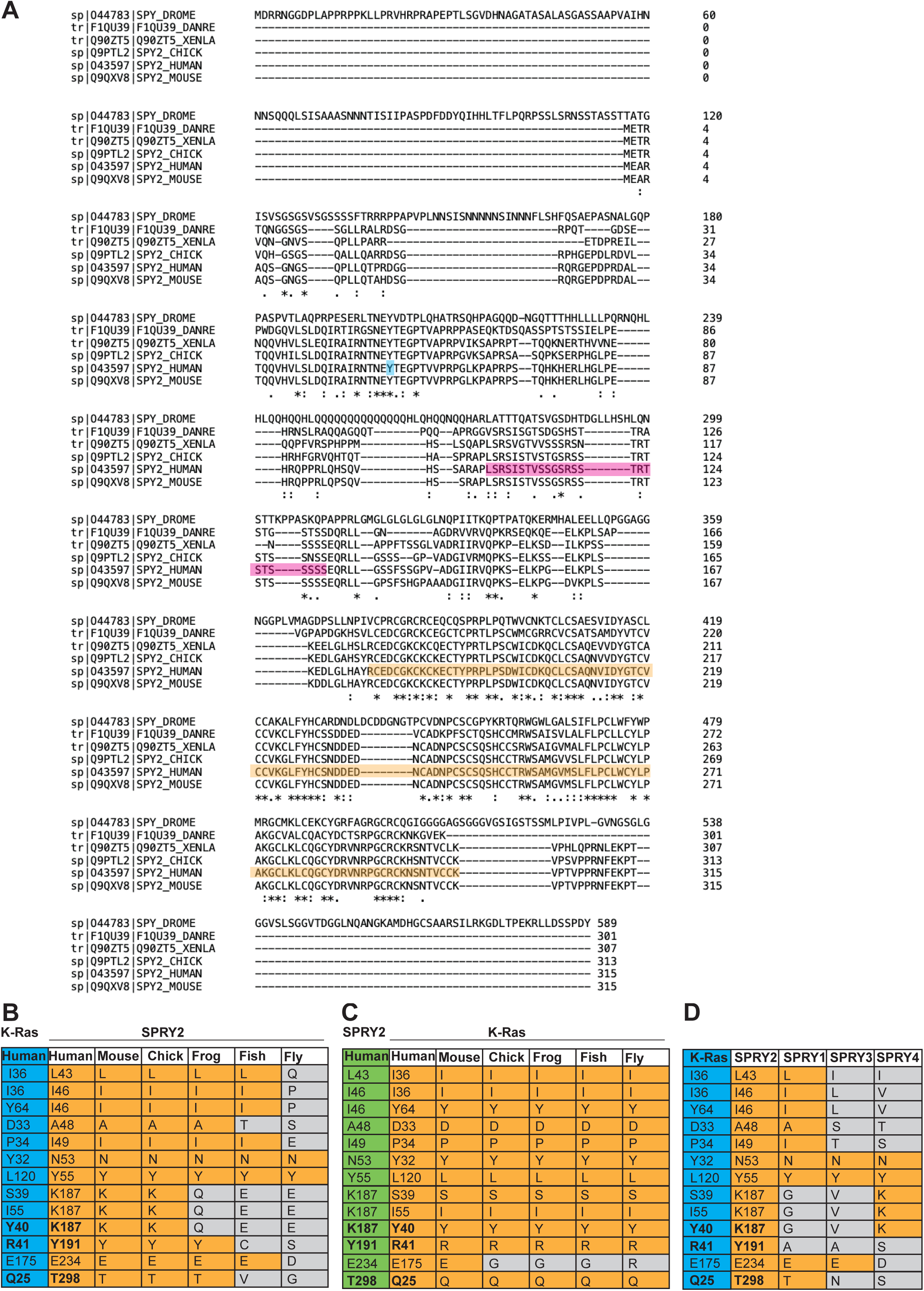
Related to main Figure 5. (**A**) Multiple sequence alignment of SPRY2 protein sequences obtained from UniProt across several species using Clustal Omega (v1.2.4). Residues of the Cysteine rich domain (orange), serine rich motif (pink) and Tyr55 (blue) at the centre of the Cbl-TKB binding motif are highlighted in the sequence of human SPRY2. Protein IDs for SPRY2 are human-O43597, mouse-Q9QXV8, chick-Q9PTL2, frog-Q90ZT5, fish-F1QU39, fly-O44783. (**B,C**) Comparison for SPRY2 (B) and K-Ras (C) of the amino acid conservation across species of all K-Ras/ SPRY2 interface residues predicted by AlphaFold3 to have a distance below 3 Å. Protein IDs for SPRY2 are given in (A). Protein IDs for K-Ras are human-P01116, mouse-P32883, chick-A0A8V0YNE2, frog-Q05147, fish-Q6AZA4, fly-P08646. (**D**) Comparison of the amino acid sequence conservation across human SPRY paralogs of residues as in (B,C). Protein IDs are SPRY2-O43597, SPRY1-O43609, SPRY3-O43610 and SPRY4-Q9C004.

### Supplementary Data (Excel File)

**Supplementary Data S1.**

Sheets1 and 2 list gene names of hits from K-Ras proximal proteome (experiment 1) and G12Ci sensitive proteome (experiment 2) experiments. Mean log2 ratios of biotinylated/non-biotinylated (for proximity) or biotinylated+G12Ci/biotinylated (for G12Ci-sensitive) from N = 3 independent biological repeats. Sheet 3 shows common hits from our meta-analysis of other published Ras proximal proteome studies. Identification of a protein with provided gene ID is annotated with a 1 if detected or 0 if not detected.

### Supplementary Table

**Supplementary Table S1.**
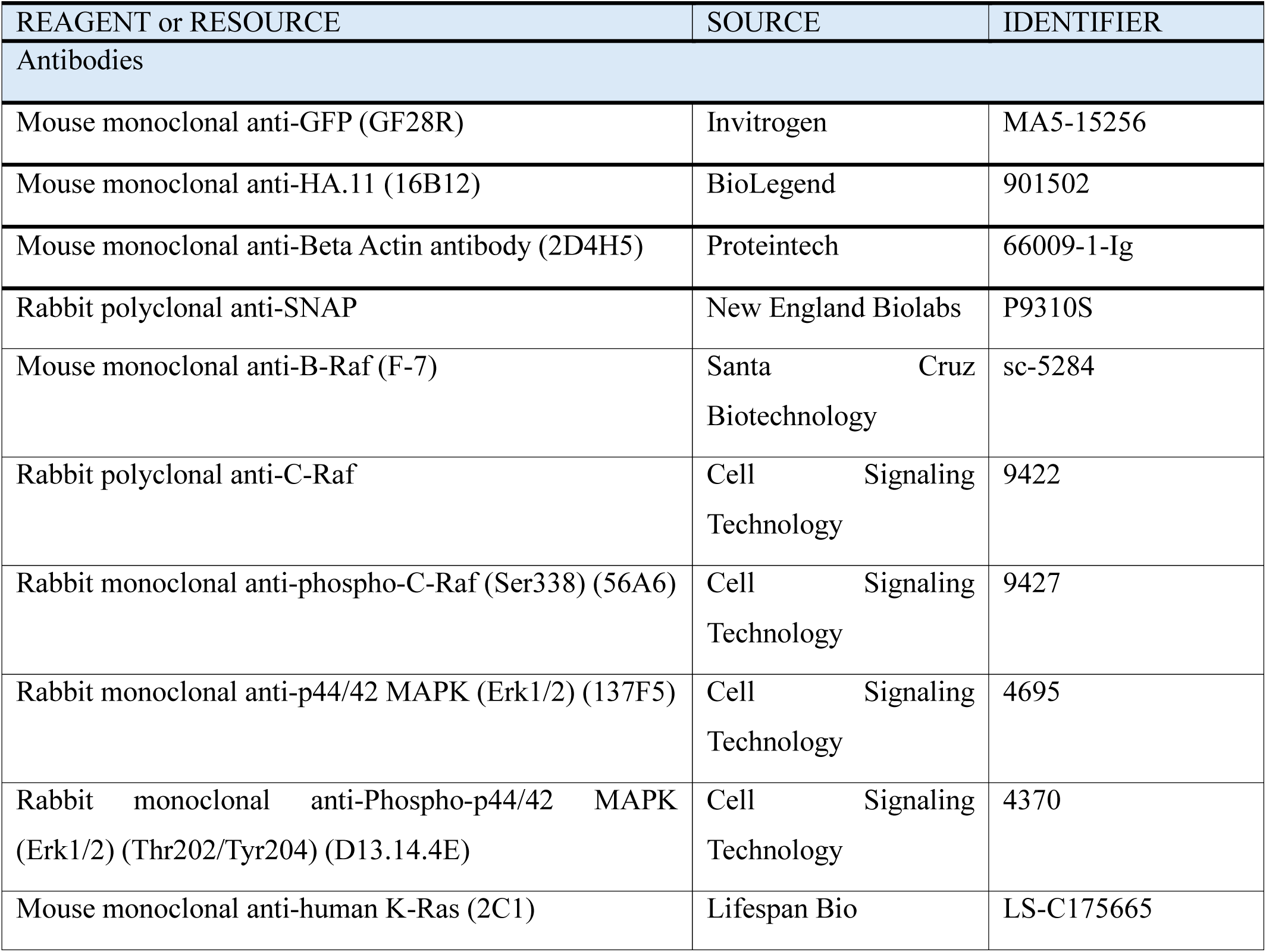
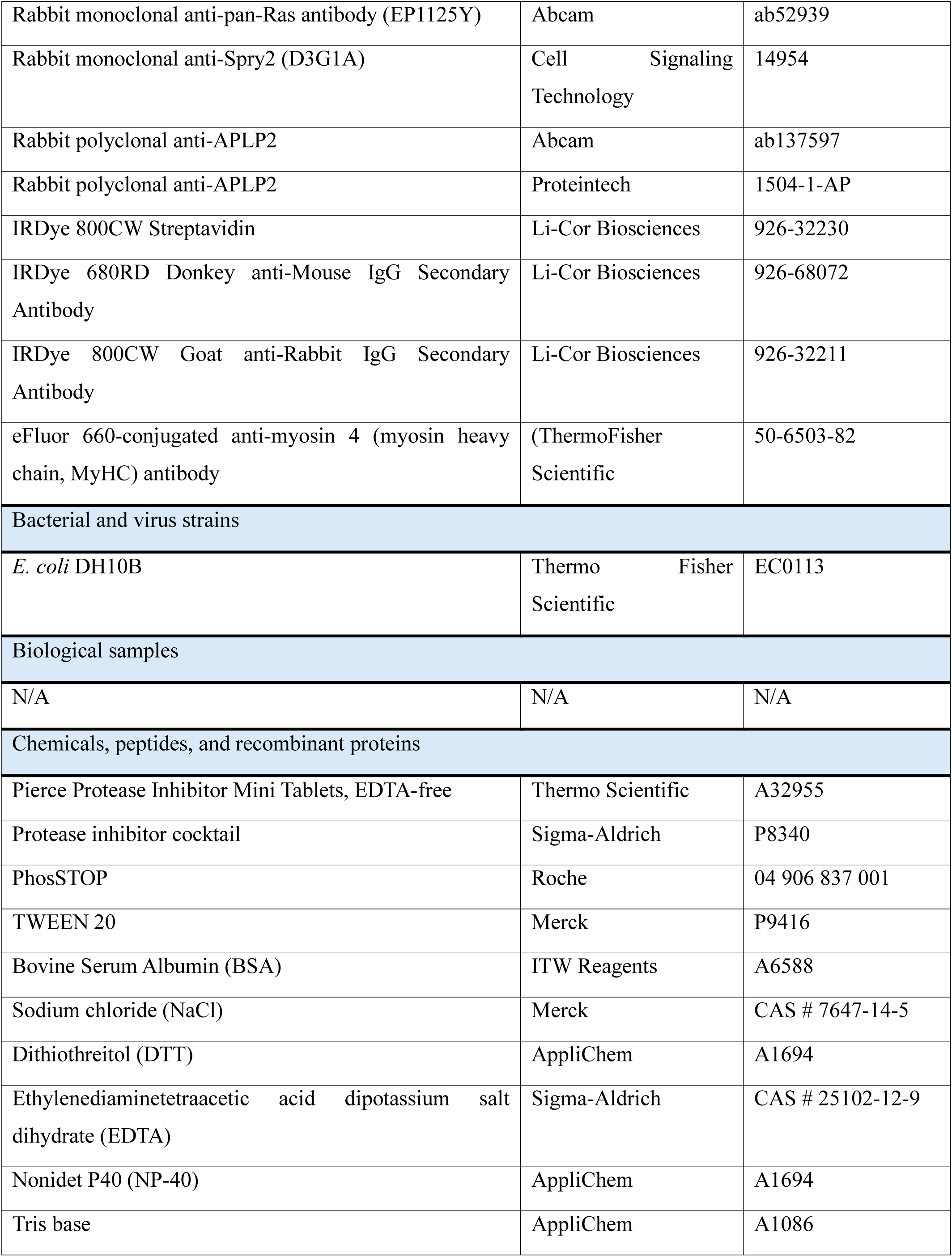
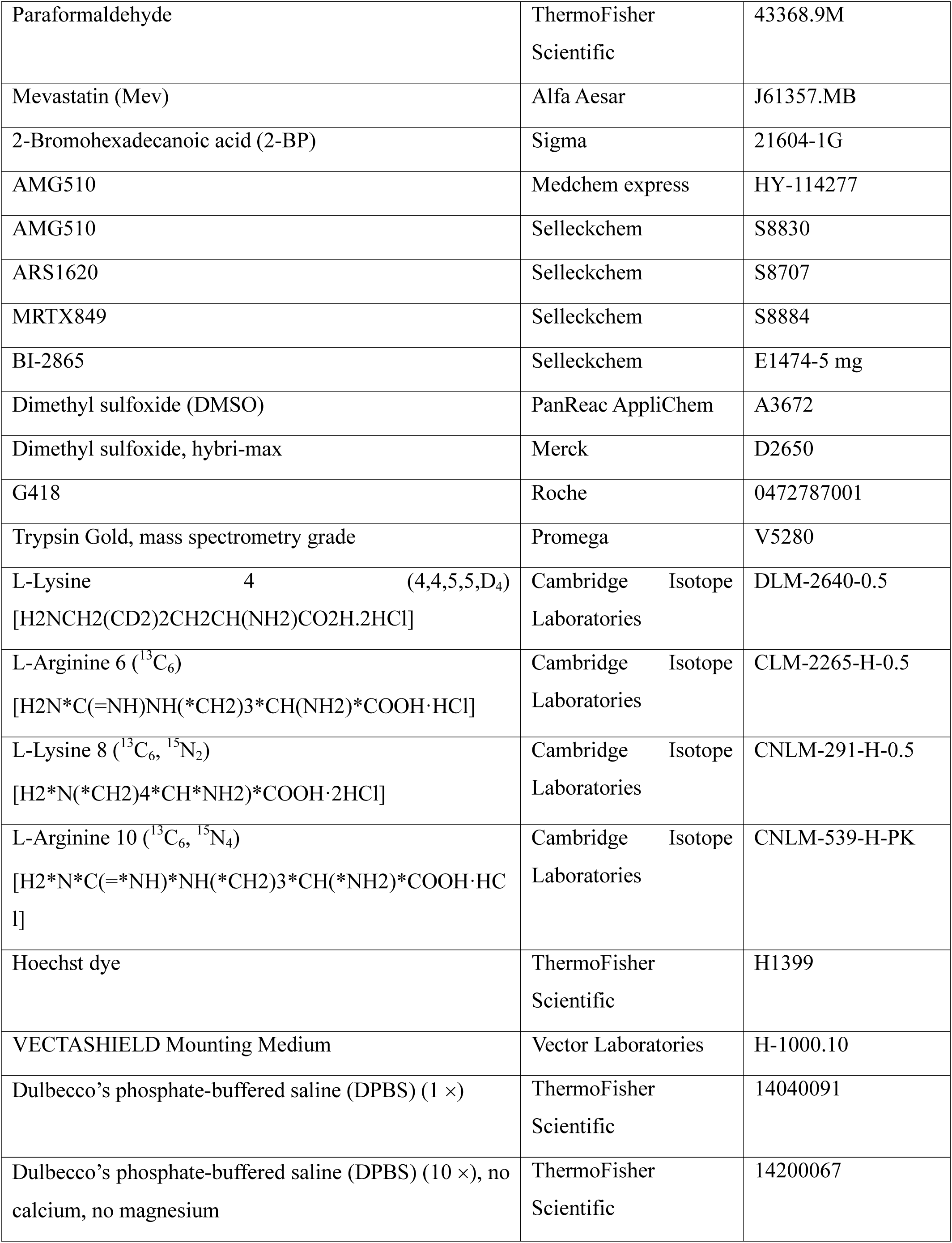
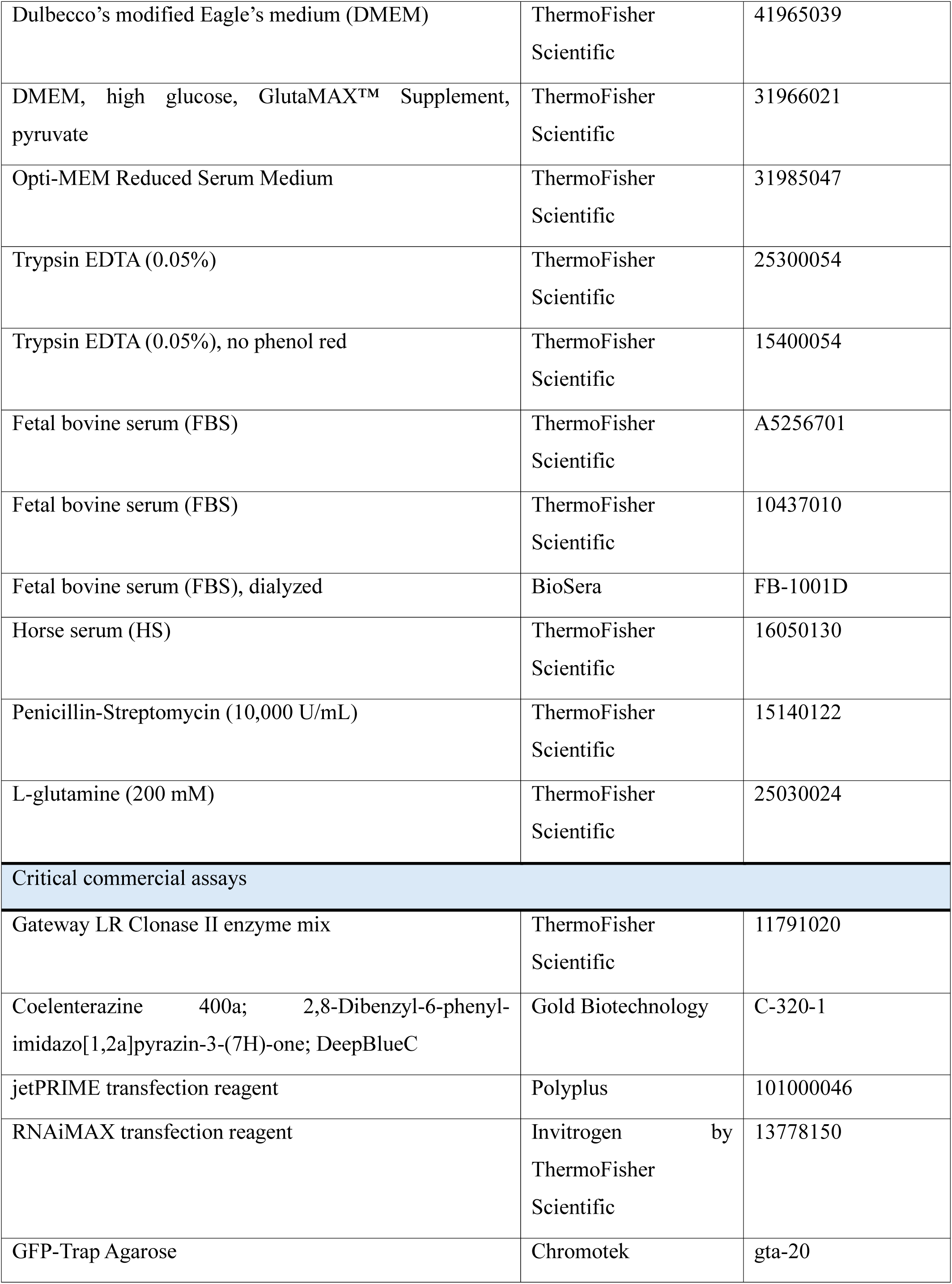
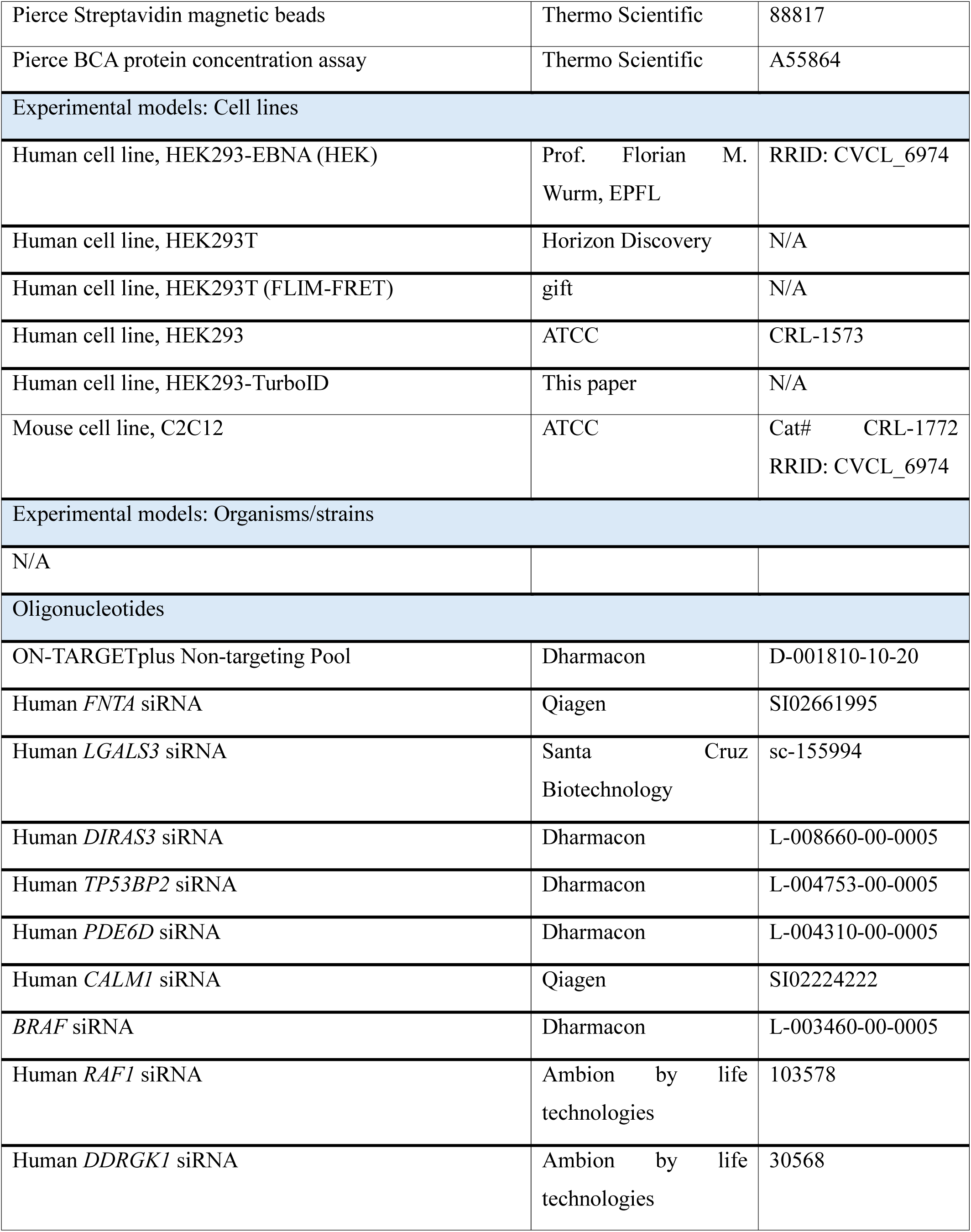
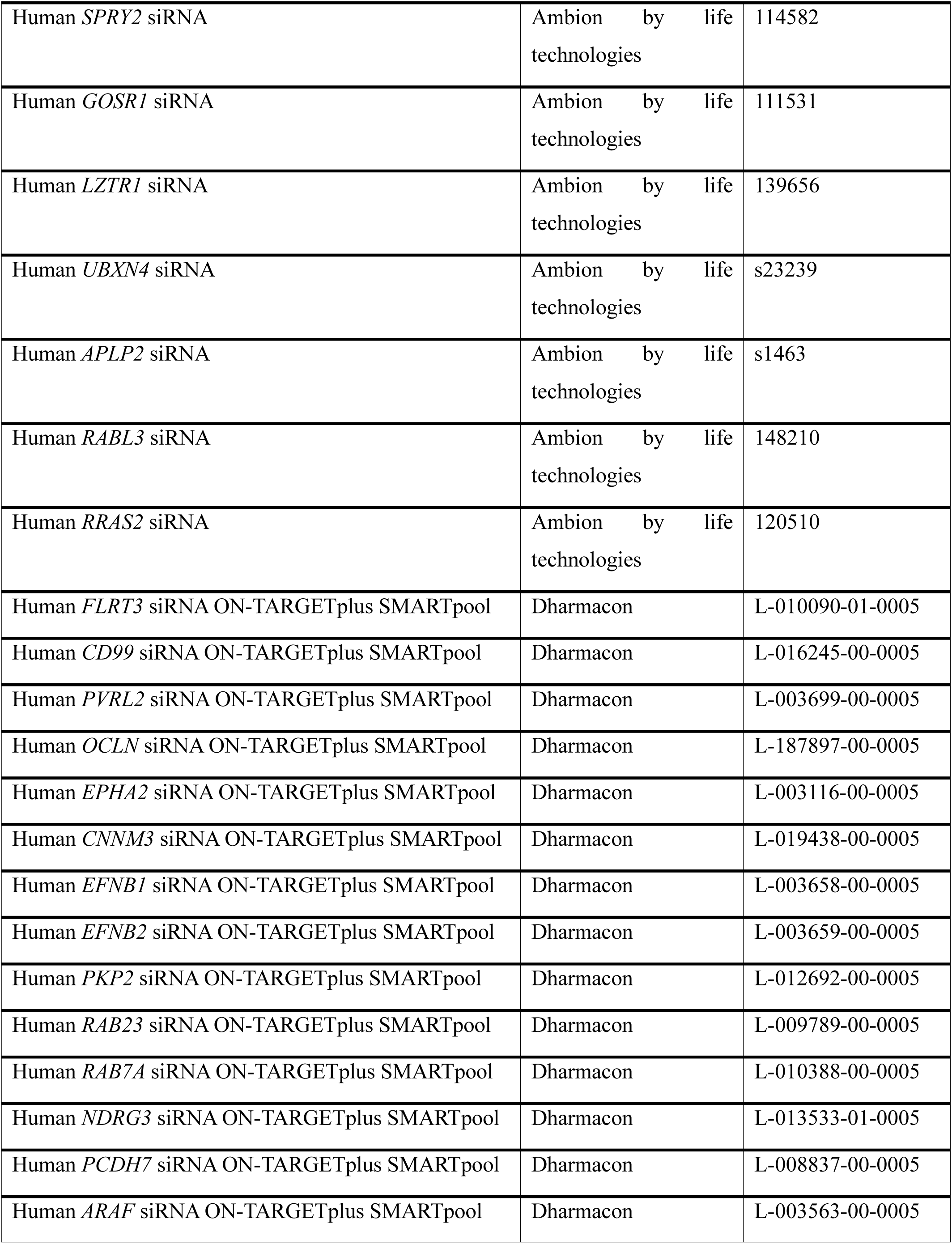
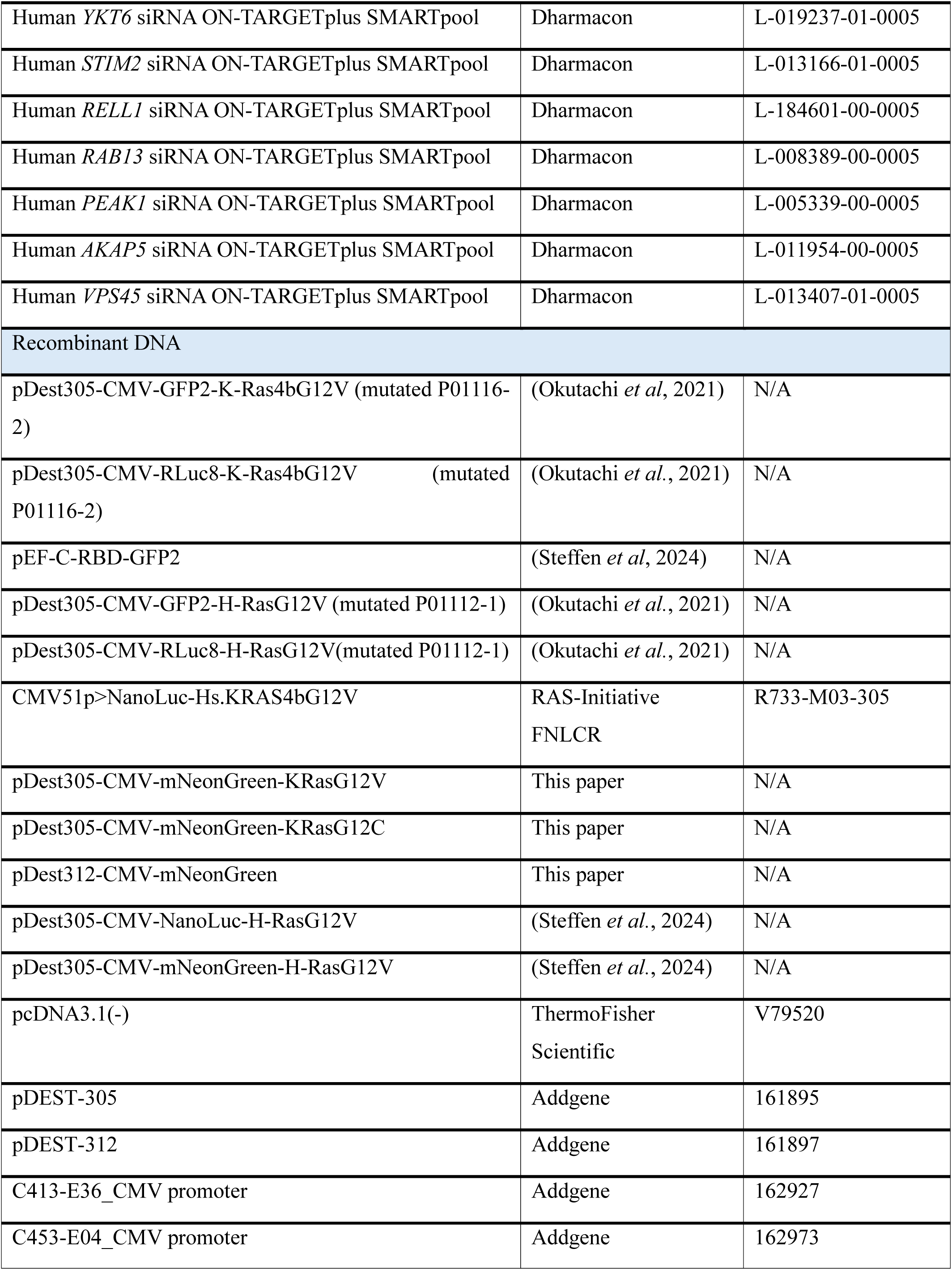
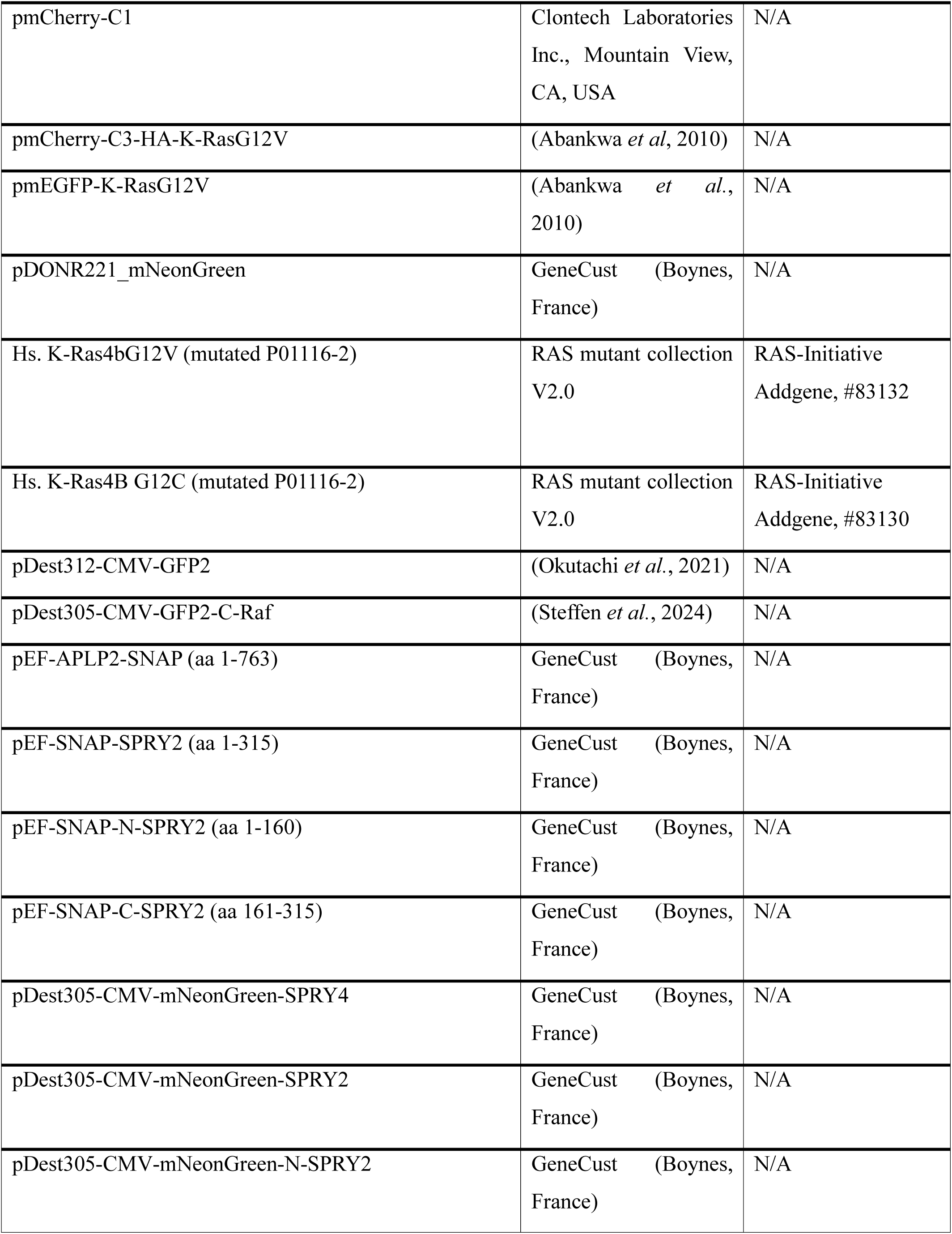
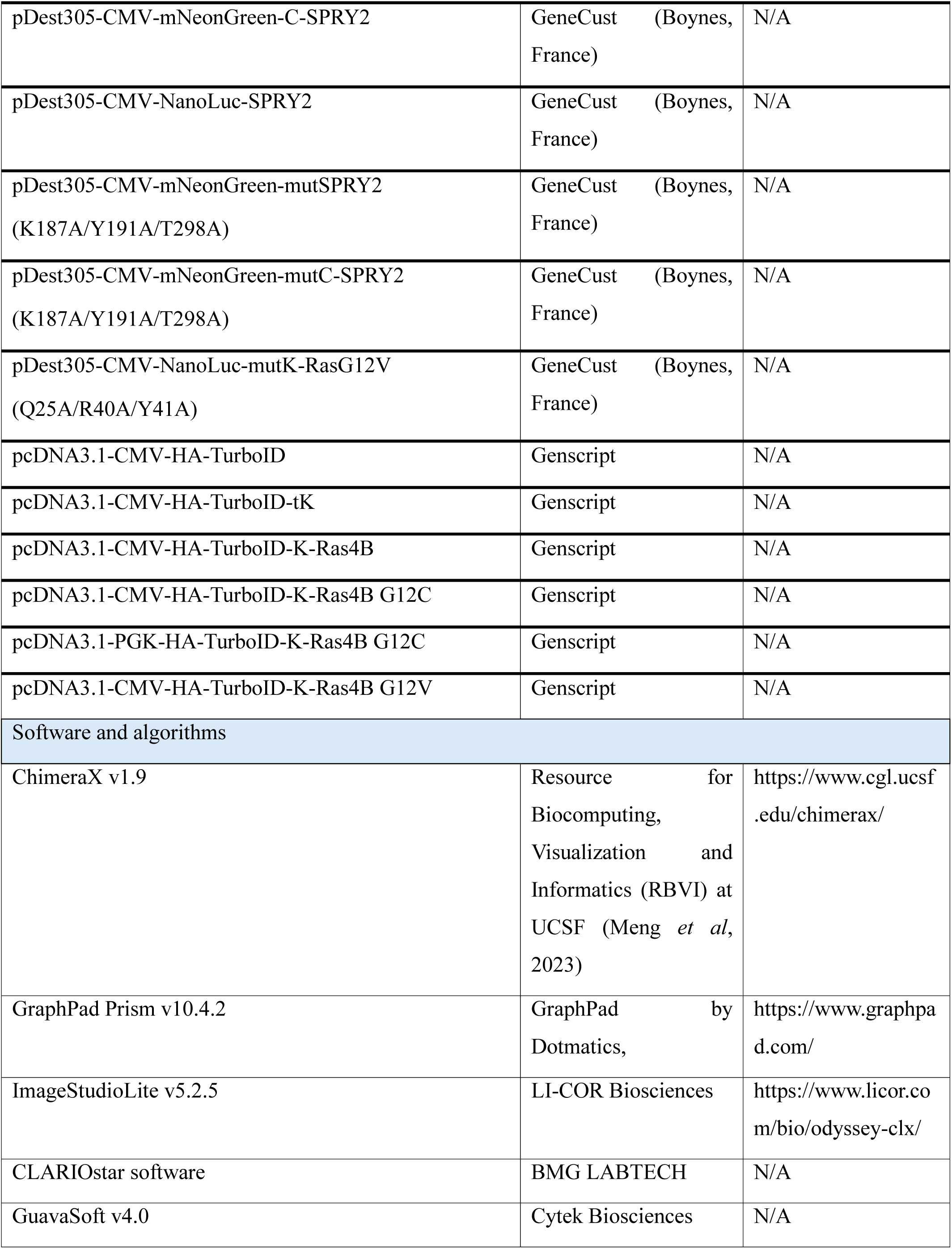
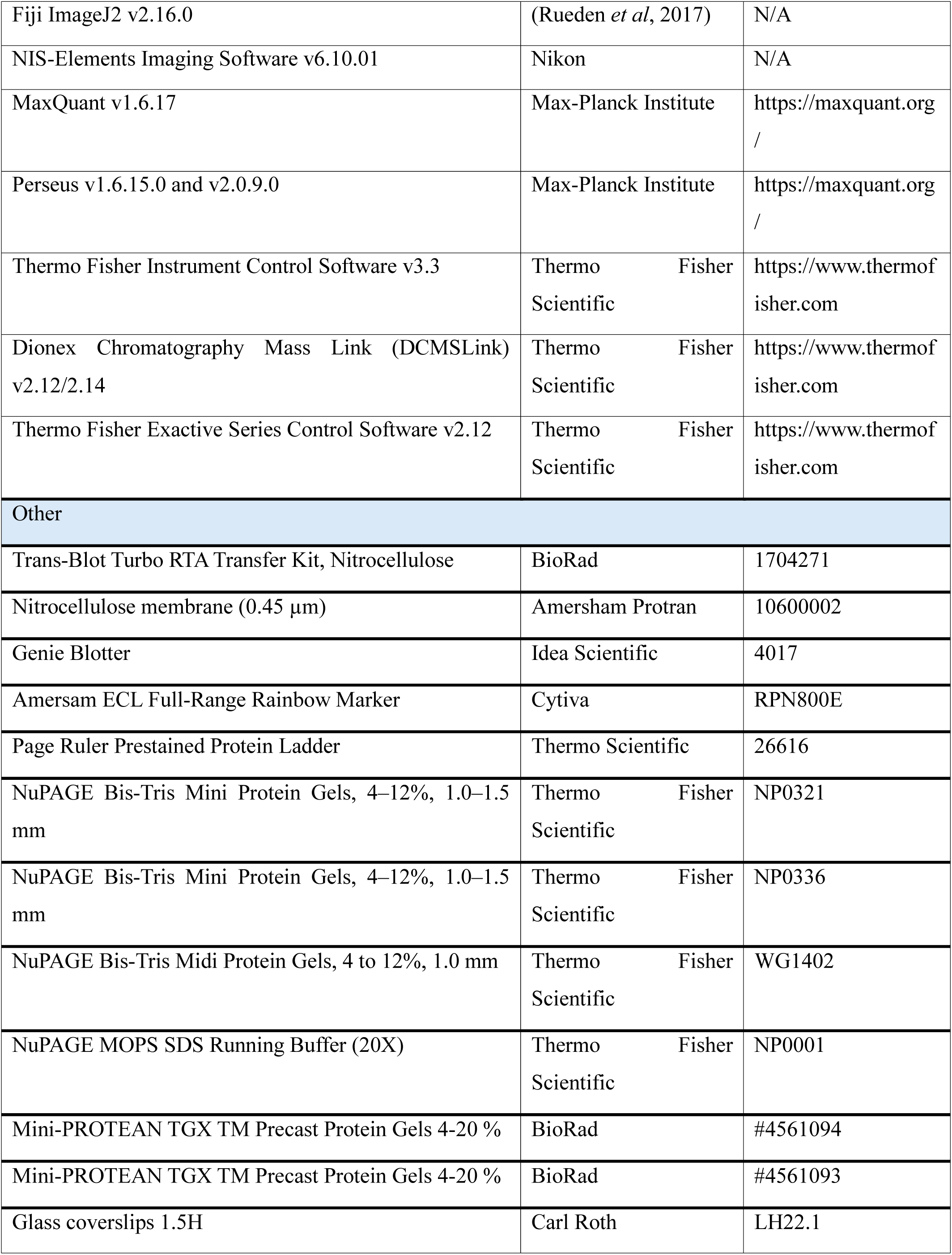
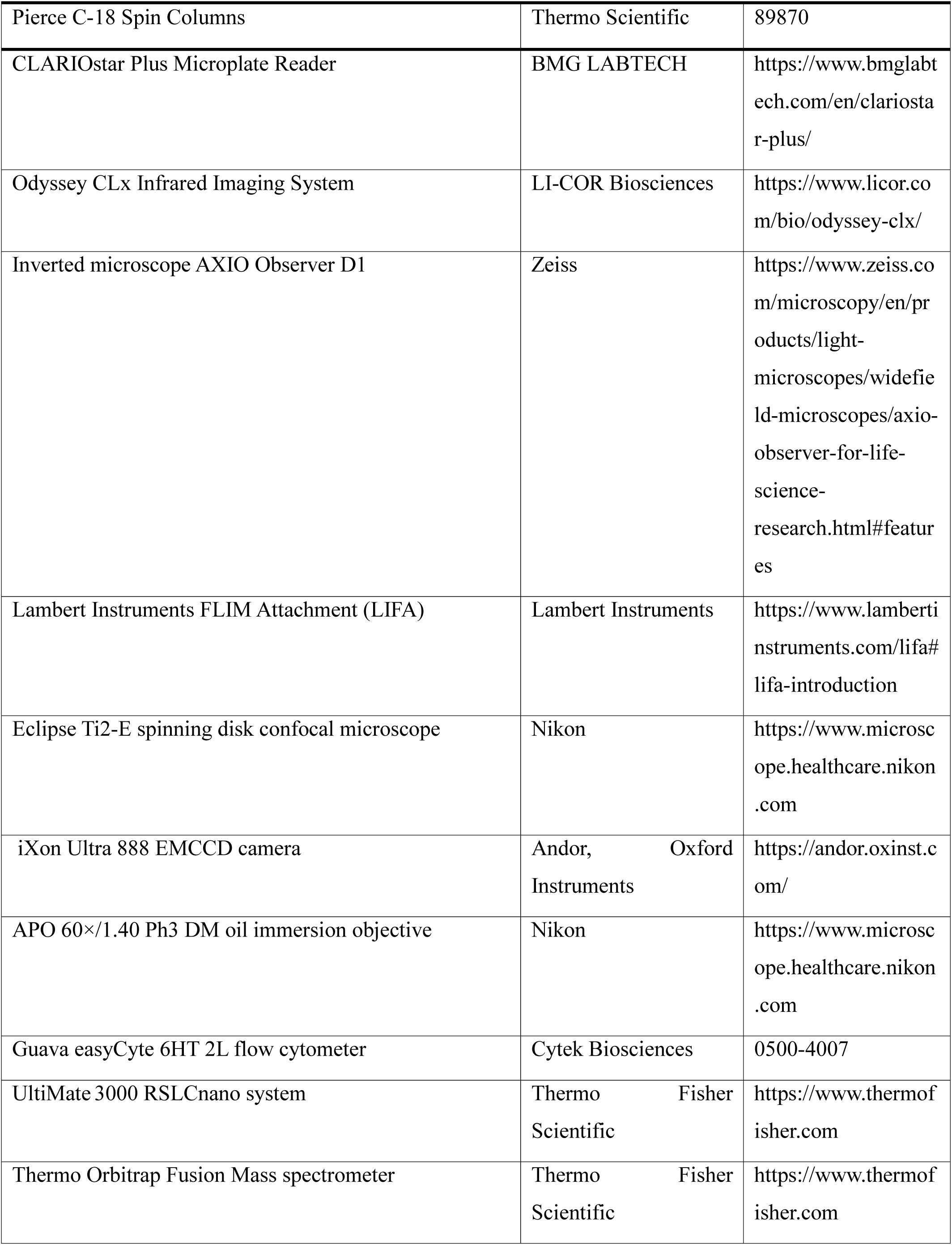
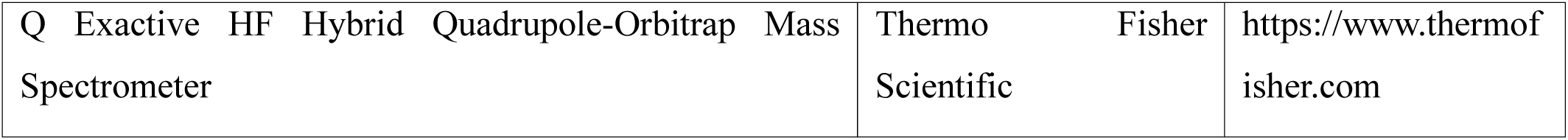
Materials, reagents, equipment and software used in this study.

## References

1. Abankwa D, Gorfe AA (2020) Mechanisms of Ras Membrane Organization and Signaling: Ras Rocks Again. Biomolecules 10

2. Abramson J, Adler J, Dunger J, Evans R, Green T, Pritzel A, Ronneberger O, Willmore L, Ballard AJ, Bambrick J et al (2024) Accurate structure prediction of biomolecular interactions with AlphaFold 3. Nature 630: 493–500

3. Adhikari H, Counter CM (2018) Interrogating the protein interactomes of RAS isoforms identifies PIP5K1A as a KRAS-specific vulnerability. Nat Commun 9: 3646

4. Akasaka K, Tamada M, Wang F, Kariya K, Shima F, Kikuchi A, Yamamoto M, Shirouzu M, Yokoyama S, Kataoka T (1996) Differential structural requirements for interaction of Ras protein with its distinct downstream effectors. J Biol Chem 271: 5353–5360

5. Barruet E, Garcia SM, Striedinger K, Wu J, Lee S, Byrnes L, Wong A, Xuefeng S, Tamaki S, Brack AS et al (2020) Functionally heterogeneous human satellite cells identified by single cell RNA sequencing. Elife 9

6. Beganton B, Coyaud E, Laurent EMN, Mange A, Jacquemetton J, Le Romancer M, Raught B, Solassol J (2020) Proximal Protein Interaction Landscape of RAS Paralogs. Cancers (Basel) 12

7. Bigot A, Duddy WJ, Ouandaogo ZG, Negroni E, Mariot V, Ghimbovschi S, Harmon B, Wielgosik A, Loiseau C, Devaney J et al (2015) Age-Associated Methylation Suppresses SPRY1, Leading to a Failure of Re-quiescence and Loss of the Reserve Stem Cell Pool in Elderly Muscle. Cell Rep 13: 1172–1182

8. Blazevits O, Mideksa YG, Solman M, Ligabue A, Ariotti N, Nakhaeizadeh H, Fansa EK, Papageorgiou AC, Wittinghofer A, Ahmadian MR et al (2016) Galectin-1 dimers can scaffold Raf-effectors to increase H-ras nanoclustering. Sci Rep 6: 24165

9. Branon TC, Bosch JA, Sanchez AD, Udeshi ND, Svinkina T, Carr SA, Feldman JL, Perrimon N, Ting AY (2018) Efficient proximity labeling in living cells and organisms with TurboID. Nat Biotechnol 36: 880–887

10. Cheng DK, Oni TE, Thalappillil JS, Park Y, Ting HC, Alagesan B, Prasad NV, Addison K, Rivera KD, Pappin DJ et al (2021) Oncogenic KRAS engages an RSK1/NF1 pathway to inhibit wild-type RAS signaling in pancreatic cancer. Proc Natl Acad Sci U S A 118

11. Chippalkatti R, Parisi B, Kouzi F, Laurini C, Ben Fredj N, Abankwa DK (2024) RAS isoform specific activities are disrupted by disease associated mutations during cell differentiation. Eur J Cell Biol 103: 151425

12. Cho KF, Branon TC, Udeshi ND, Myers SA, Carr SA, Ting AY (2020) Proximity labeling in mammalian cells with TurboID and split-TurboID. Nat Protoc 15: 3971–3999

13. de Alvaro C, Martinez N, Rojas JM, Lorenzo M (2005) Sprouty-2 overexpression in C2C12 cells confers myogenic differentiation properties in the presence of FGF2. Mol Biol Cell 16: 4454–4461

14. Duval CJ, Steffen CL, Pavic K, Abankwa DK (2024) Protocol to measure and analyze protein interactions in mammalian cells using bioluminescence resonance energy transfer. STAR Protoc 5: 103348

15. Egan JE, Hall AB, Yatsula BA, Bar-Sagi D (2002) The bimodal regulation of epidermal growth factor signaling by human Sprouty proteins. Proc Natl Acad Sci U S A 99: 6041–6046

16. Go CD, Knight JDR, Rajasekharan A, Rathod B, Hesketh GG, Abe KT, Youn JY, Samavarchi-Tehrani P, Zhang H, Zhu LY et al (2021) A proximity-dependent biotinylation map of a human cell. Nature 595: 120–124

17. Goldfinger LE, Ptak C, Jeffery ED, Shabanowitz J, Han J, Haling JR, Sherman NE, Fox JW, Hunt DF, Ginsberg MH (2007) An experimentally derived database of candidate Ras-interacting proteins. J Proteome Res 6: 1806–1811

18. Gross I, Bassit B, Benezra M, Licht JD (2001) Mammalian sprouty proteins inhibit cell growth and differentiation by preventing ras activation. J Biol Chem 276: 46460–46468

19. Guy GR, Jackson RA, Yusoff P, Chow SY (2009) Sprouty proteins: modified modulators, matchmakers or missing links? Journal of Endocrinology 203: 191–202

20. Guzman C, Oetken-Lindholm C, Abankwa D (2016) Automated High-Throughput Fluorescence Lifetime Imaging Microscopy to Detect Protein-Protein Interactions. J Lab Autom 21: 238–245

21. Guzman C, Solman M, Ligabue A, Blazevits O, Andrade DM, Reymond L, Eggeling C, Abankwa D (2014) The efficacy of Raf kinase recruitment to the GTPase H-ras depends on H-ras membrane conformer-specific nanoclustering. J Biol Chem 289: 9519–9533

22. Hacohen N, Kramer S, Sutherland D, Hiromi Y, Krasnow MA (1998) sprouty encodes a novel antagonist of FGF signaling that patterns apical branching of the Drosophila airways. Cell 92: 253–263

23. Hanafusa H, Torii S, Yasunaga T, Nishida E (2002) Sprouty1 and Sprouty2 provide a control mechanism for the Ras/MAPK signalling pathway. Nat Cell Biol 4: 850–858

24. Hedberg C, Dekker FJ, Rusch M, Renner S, Wetzel S, Vartak N, Gerding-Reimers C, Bon RS, Bastiaens PI, Waldmann H (2011) Development of highly potent inhibitors of the Ras-targeting human acyl protein thioesterases based on substrate similarity design. Angew Chem Int Ed Engl 50: 9832–9837

25. Hobbs GA, Der CJ, Rossman KL (2016) RAS isoforms and mutations in cancer at a glance. J Cell Sci 129: 1287–1292

26. Jung JE, Moon SH, Kim DK, Choi C, Song J, Park KS (2012) Sprouty1 regulates neural and endothelial differentiation of mouse embryonic stem cells. Stem Cells Dev 21: 554–561

27. Kaya P, Schaffner-Reckinger E, Manoharan GB, Vukic V, Kiriazis A, Ledda M, Burgos Renedo M, Pavic K, Gaigneaux A, Glaab E et al (2024) An Improved PDE6D Inhibitor Combines with Sildenafil To Inhibit KRAS Mutant Cancer Cell Growth. J Med Chem 67: 8569–8584

28. Kim D, Herdeis L, Rudolph D, Zhao Y, Bottcher J, Vides A, Ayala-Santos CI, Pourfarjam Y, Cuevas-Navarro A, Xue JY et al (2023) Pan-KRAS inhibitor disables oncogenic signalling and tumour growth. Nature 619: 160–166

29. Kovalski JR, Bhaduri A, Zehnder AM, Neela PH, Che Y, Wozniak GG, Khavari PA (2019) The Functional Proximal Proteome of Oncogenic Ras Includes mTORC2. Mol Cell 73: 830–844 e812

30. Lim J, Yusoff P, Wong ES, Chandramouli S, Lao DH, Fong CW, Guy GR (2002) The cysteine-rich sprouty translocation domain targets mitogen-activated protein kinase inhibitory proteins to phosphatidylinositol 4,5-bisphosphate in plasma membranes. Mol Cell Biol 22: 7953–7966

31. Lito P, Mets BD, Kleff S, O’Reilly S, Maher VM, McCormick JJ (2008) Evidence that sprouty 2 is necessary for sarcoma formation by H-Ras oncogene-transformed human fibroblasts. J Biol Chem 283: 2002–2009

32. Locatelli C, Lemonidis K, Salaun C, Tomkinson NCO, Chamberlain LH (2020) Identification of key features required for efficient S-acylation and plasma membrane targeting of sprouty-2. J Cell Sci 133

33. Lu H, Shi X, Wu G, Zhu J, Song C, Zhang Q, Yang G (2015) FGF13 regulates proliferation and differentiation of skeletal muscle by down-regulating Spry1. Cell Prolif 48: 550–560

34. Manoharan GB, Laurini C, Bottone S, Ben Fredj N, Abankwa DK (2023) K-Ras Binds Calmodulin-Related Centrin1 with Potential Implications for K-Ras Driven Cancer Cell Stemness. Cancers (Basel) 15

35. Manoharan GB, Okutachi S, Abankwa D (2022) Potential of phenothiazines to synergistically block calmodulin and reactivate PP2A in cancer cells. PLoS One 17: e0268635

36. Masoumi-Moghaddam S, Amini A, Morris DL (2014) The developing story of Sprouty and cancer. Cancer Metastasis Rev 33: 695–720

37. Ng C, Jackson RA, Buschdorf JP, Sun Q, Guy GR, Sivaraman J (2008) Structural basis for a novel intrapeptidyl H-bond and reverse binding of c-Cbl-TKB domain substrates. EMBO J 27: 804–816

38. Okutachi S, Manoharan GB, Kiriazis A, Laurini C, Catillon M, McCormick F, Yli-Kauhaluoma J, Abankwa D (2021) A Covalent Calmodulin Inhibitor as a Tool to Study Cellular Mechanisms of K-Ras-Driven Stemness. Front Cell Dev Biol 9: 665673

39. Ong SE, Blagoev B, Kratchmarova I, Kristensen DB, Steen H, Pandey A, Mann M (2002) Stable isotope labeling by amino acids in cell culture, SILAC, as a simple and accurate approach to expression proteomics. Mol Cell Proteomics 1: 376–386

40. Ozaki K, Miyazaki S, Tanimura S, Kohno M (2005) Efficient suppression of FGF-2-induced ERK activation by the cooperative interaction among mammalian Sprouty isoforms. J Cell Sci 118: 5861–5871

41. Pandey P, Sliker B, Peters HL, Tuli A, Herskovitz J, Smits K, Purohit A, Singh RK, Dong J, Batra SK et al (2016) Amyloid precursor protein and amyloid precursor-like protein 2 in cancer. Oncotarget 7: 19430–19444

42. Parisi B, Sünnen M, Chippalkatti R, Abankwa DK (2023) A flow-cytometry-based pipeline for the rapid quantification of C2C12 cell differentiation. STAR Protoc: 102637

43. Parkkola H, Siddiqui FA, Oetken-Lindholm C, Abankwa D (2021) FLIM-FRET Analysis of Ras Nanoclustering and Membrane-Anchorage. Methods Mol Biol 2262: 233–250

44. Pavic K, Chippalkatti R, Abankwa D (2022) Drug targeting opportunities en route to Ras nanoclusters. Adv Cancer Res 153: 63–99

45. Phoenix TN, Temple S (2010) Spred1, a negative regulator of Ras-MAPK-ERK, is enriched in CNS germinal zones, dampens NSC proliferation, and maintains ventricular zone structure. Genes Dev 24: 45–56

46. Poelaert BJ, Knoche SM, Larson AC, Pandey P, Seshacharyulu P, Khan N, Maurer HC, Olive KP, Sheinin Y, Ahmad R et al (2021) Amyloid Precursor-like Protein 2 Expression Increases during Pancreatic Cancer Development and Shortens the Survival of a Spontaneous Mouse Model of Pancreatic Cancer. Cancers (Basel) 13

47. Posada IM, Serulla M, Zhou Y, Oetken-Lindholm C, Abankwa D, Lectez B (2016) ASPP2 Is a Novel Pan-Ras Nanocluster Scaffold. PLoS One 11: e0159677

48. Prior IA, Harding A, Yan J, Sluimer J, Parton RG, Hancock JF (2001) GTP-dependent segregation of H-ras from lipid rafts is required for biological activity. Nat Cell Biol 3: 368–375

49. Prior IA, Hood FE, Hartley JL (2020) The Frequency of Ras Mutations in Cancer. Cancer Res 80: 2969–2974

50. Puranik N, Jung H, Song M (2024) SPROUTY2, a Negative Feedback Regulator of Receptor Tyrosine Kinase Signaling, Associated with Neurodevelopmental Disorders: Current Knowledge and Future Perspectives. Int J Mol Sci 25

51. Rolland T, Tasan M, Charloteaux B, Pevzner SJ, Zhong Q, Sahni N, Yi S, Lemmens I, Fontanillo C, Mosca R et al (2014) A proteome-scale map of the human interactome network. Cell 159: 1212–1226

52. Roux KJ, Kim DI, Raida M, Burke B (2012) A promiscuous biotin ligase fusion protein identifies proximal and interacting proteins in mammalian cells. J Cell Biol 196: 801–810

53. Sasaki A, Taketomi T, Kato R, Saeki K, Nonami A, Sasaki M, Kuriyama M, Saito N, Shibuya M, Yoshimura A (2003) Mammalian Sprouty4 suppresses Ras-independent ERK activation by binding to Raf1. Nat Cell Biol 5: 427–432

54. Shea KL, Xiang W, LaPorta VS, Licht JD, Keller C, Basson MA, Brack AS (2010) Sprouty1 regulates reversible quiescence of a self-renewing adult muscle stem cell pool during regeneration. Cell Stem Cell 6: 117–129

55. Siddiqui FA, Alam C, Rosenqvist P, Ora M, Sabt A, Manoharan GB, Bindu L, Okutachi S, Catillon M, Taylor T et al (2020) PDE6D Inhibitors with a New Design Principle Selectively Block K-Ras Activity. ACS Omega 5: 832–842

56. Siddiqui FA, Parkkola H, Vukic V, Oetken-Lindholm C, Jaiswal A, Kiriazis A, Pavic K, Aittokallio T, Salminen TA, Abankwa D (2021) Novel Small Molecule Hsp90/Cdc37 Interface Inhibitors Indirectly Target K-Ras-Signaling. Cancers (Basel) 13

57. Simanshu DK, Nissley DV, McCormick F (2017) RAS Proteins and Their Regulators in Human Disease. Cell 170: 17–33

58. Smith MJ (2023) Defining bone fide effectors of RAS GTPases. Bioessays 45: e2300088

59. Song J, Wang B, Zou M, Zhou H, Ding Y, Ren W, Fang L, Zhang J (2025) Mapping the Interactome of KRAS and Its G12C/D/V Mutants by Integrating TurboID Proximity Labeling with Quantitative Proteomics. Biology (Basel) 14

60. Steffen CL, Manoharan GB, Pavic K, Yeste-Vazquez A, Knuuttila M, Arora N, Zhou Y, Harma H, Gaigneaux A, Grossmann TN et al (2024) Identification of an H-Ras nanocluster disrupting peptide. Commun Biol 7: 837

61. Sun Q, Jackson RA, Ng C, Guy GR, Sivaraman J (2010) Additional serine/threonine phosphorylation reduces binding affinity but preserves interface topography of substrate proteins to the c-Cbl TKB domain. PLoS One 5: e12819

62. Wakioka T, Sasaki A, Kato R, Shouda T, Matsumoto A, Miyoshi K, Tsuneoka M, Komiya S, Baron R, Yoshimura A (2001) Spred is a Sprouty-related suppressor of Ras signalling. Nature 412: 647–651

63. Wall VE, Garvey LA, Mehalko JL, Procter LV, Esposito D (2014) Combinatorial assembly of clone libraries using site-specific recombination. Methods Mol Biol 1116: 193–208

## Supplementary Information References

65. Abankwa D, Gorfe AA, Inder K, Hancock JF (2010) Ras membrane orientation and nanodomain localization generate isoform diversity. Proc Natl Acad Sci U S A 107: 1130–1135

66. Meng EC, Goddard TD, Pettersen EF, Couch GS, Pearson ZJ, Morris JH, Ferrin TE (2023) UCSF ChimeraX: Tools for structure building and analysis. Protein Sci 32: e4792

67. Okutachi S, Manoharan GB, Kiriazis A, Laurini C, Catillon M, McCormick F, Yli-Kauhaluoma J, Abankwa D (2021) A Covalent Calmodulin Inhibitor as a Tool to Study Cellular Mechanisms of K-Ras-Driven Stemness. Front Cell Dev Biol 9: 665673

68. Rueden CT, Schindelin J, Hiner MC, DeZonia BE, Walter AE, Arena ET, Eliceiri KW (2017) ImageJ2: ImageJ for the next generation of scientific image data. BMC Bioinformatics 18: 529

69. Steffen CL, Manoharan GB, Pavic K, Yeste-Vazquez A, Knuuttila M, Arora N, Zhou Y, Harma H, Gaigneaux A, Grossmann TN et al (2024) Identification of an H-Ras nanocluster disrupting peptide. Commun Biol 7: 837

